# The genetic and biochemical determinants of mRNA degradation rates in mammals

**DOI:** 10.1101/2022.03.18.484474

**Authors:** Vikram Agarwal, David Kelley

**Affiliations:** Calico Life Sciences LLC, South San Francisco, CA 94080, USA

**Keywords:** mRNA stability, mRNA half-life, deep neural networks, post-transcriptional gene regulation

## Abstract

**Background:** Degradation rate is a fundamental aspect of mRNA metabolism, and the factors governing it remain poorly characterized. Understanding the genetic and biochemical determinants of mRNA half-life would enable a more precise identification of variants that perturb gene expression through post-transcriptional gene regulatory mechanisms.

**Results:** Here, we establish a compendium of 54 human and 27 mouse transcriptome-wide mRNA decay rate datasets. A meta-analysis of these data identified a prevalence of technical noise and measurement bias, induced partially by the underlying experimental strategy. Correcting for these biases allowed us to derive more precise, consensus measurements of half-life which exhibit enhanced consistency between species. We trained substantially improved statistical models based upon genetic and biochemical features to better predict half-life and characterize the factors molding it. Our state-of-the-art model, Saluki, is a hybrid convolutional and recurrent deep neural network which relies only upon an mRNA sequence annotated with coding frame and splice sites to predict half-life (r=0.77). Saluki predicts the impact of RNA sequences and genetic mutations therein on mRNA stability, in agreement with functional measurements derived from massively parallel reporter assays.

**Conclusions:** Our work produces a more robust “ground truth” with regards to transcriptome-wide mRNA half-lives in mammalian cells. Using these consolidated measurements, we trained a model that is over 50% more accurate in predicting half-life from sequence than existing models. Our best model, Saluki, succinctly captures many of the known determinants of mRNA half-life and can be rapidly deployed to predict the functional consequences of arbitrary mutations in the transcriptome.

## INTRODUCTION

The steady-state level of RNA is governed by two opposing forces: the rate of transcription and the rate of decay. While much headway has been made in the problem of predicting steady-state mRNA abundances through the lens of DNA-encoded features that influence transcription rates [1–5], relatively less is known about the RNA-encoded determinants that govern decay rates. Models to predict half-life from RNA sequence in mammals (achieving r^2^=0.20 and r^2^=0.39) [6, 7] have severely lagged behind the performance of those in yeast (achieving r^2^=0.59) [8]. Integrating the modeling of transcription and RNA decay into a unified model to predict steady-state mRNA levels [2] from genetic sequences would elucidate the full spectrum of regulatory functions of non-coding DNA. Consequently, this would advance the goals of providing a mechanistic explanation for evolutionary constraint in conserved sequences [9–11], identifying causal eQTLs [12, 13], diagnosing pathogenic non-coding genetic variants [14], and designing more stable and effective mRNA therapeutics [15].

The rate of mRNA decay is experimentally measured as its half-life, or the time elapsed until the initial RNA concentration has decreased by half. Experimental measurement of transcriptome-wide half-life can be achieved by one of two strategies: i) the application of transcriptional inhibitors [e.g., Actinomycin D (ActD) and α-Amanitin] to cells, or ii) a pulse-chase based method of pulsing modified nucleosides (e.g., 4sU, 5EU, and BrU) followed by chasing with unmodified nucleosides to distinguish newly synthesized RNA from pre-existing RNA [16]. Both strategies are followed by the profiling of RNA levels over a time course, and data for each gene are fit to an exponential decay curve to ascertain the gene’s half-life. Although it has long been appreciated that different methods induce measurement bias [16–18], many modern studies that deploy these methodologies fail to acknowledge the prevalence and impact of these biases on result interpretation.

Measurement biases emerge for a multitude of reasons. Transcriptional inhibitors are convenient, but do not necessarily enter all cells to fully block transcription [16]. Moreover, such drugs can lead to cytotoxicity [17] or impact unintended pathways such as translation, thus preventing the attachment of mRNA to ribosomes and resulting in artificially altered mRNA metabolism [16, 18]. Pulse-chase methods are inaccurate in the scenario in which the half-life is shorter than the chase period, and cellular parameters such as doubling time and drug uptake further bias half-life measurement [18]. Finally, the incorporation of uridine analogs is thought to be a stochastic process that is proportional to the number of Us in an RNA, leading to RNA-length-dependent labeling and enrichment biases [19, 20]. Collectively, the ultimate consequence of these biases is the disagreement between different methods in the physiologically relevant estimate of half-life [16].

It has been postulated that considering an ensemble of different methods would empower a more precise measurement of half-life, providing a path towards circumventing the tradeoffs and limitations among any individual method [18]. While a meta-analysis of half-life datasets in yeast revealed surprisingly discordant results among half-lives measured by different research groups [21], the consistency among half-life datasets in mammalian organisms remains largely uncharacterized. Collecting such a compendium of datasets would potentially enable the derivation of consensus measurements of cellular mRNA half-life in a fashion that is less obfuscated by technical noise and methodological bias.

A precise measurement of RNA half-life would enable a clear-eyed examination of how different molecular pathways modulate half-life relative to one another. Numerous RNA-binding proteins (RBPs) and sequence-encoded features have been implicated in regulating half-life. Examples include: i) generic features of an mRNA such as its GC content [22], length, and ORF exon junction density [6, 7]; ii) the presence of microRNA (miRNA) binding sites [23–25]; iii) codon frequencies and interactions with the translation machinery [8,26–30]; iv) mRNA structure [31, 32]; v) Pumilio binding elements [33]; vi) AU-rich elements (AREs) [16, 34]; and vii) YTHDF proteins [35] via m6A recognition [36, 37].

Attempts to examine the relative contribution of sequence and biochemical features to the specification of half-life [6–8,38–40] have been undermined by half-life measurement biases, and have not exhaustively considered all of the known pathways that affect RNA stability. In this study, we assembled a compendium of 54 human and 27 mouse mRNA half-life datasets to derive more precise, consensus measurements of RNA stability in mammalian cells. Using our enhanced measurements, we derived improved genetic and biochemical models towards the goals of quantifying the relative influence of different pathways and improving the predictability of half-life from such features. Our state-of-the-art model Saluki, a hybrid convolutional and recurrent neural network, is capable of predicting the effects of RNA sequences and genetic variants therein on RNA stability, in agreement with functional measurements derived from massively parallel reporter assays.

## RESULTS

### Comparison of study concordance in the mammalian half-life compendium

To generate a compendium of mammalian half-life datasets, we mined the literature for published human and mouse half-life data. In total, we identified 33 publications reporting transcriptome-wide half-lives, with 16 submitting human data, 14 submitting mouse data, and 3 submitting data from both species (**Table 1**). Counting individual replicates, this led to a total of 54 human and 27 mouse half-life measurements. The human studies encompassed 10 cell types and 5 measurement procedures, while the mouse studies encompassed 8 cell types and 3 procedures (**Table 1**). Each of the human studies reported half-lives for between ∼2,500-13,000 genes (**Additional file 1: Fig. S1**), while those from the mouse reported half-lives for between ∼500-18,000 genes (**Additional file 1: Fig. S2a**) (**Additional file 2: Dataset S1**).

**Table 1.** Overview of studies compiled for meta-analysis, listing each publication by a GroupID, the method of determining half-lives, the species of origin (M: mouse, H: human), the cell type of origin, and number of samples (comma-separated when listing human and mouse, respectively).

| GroupID | Method | Species | Cell types | Samples | Reference |
| --- | --- | --- | --- | --- | --- |
| Akimitsu | BrU (BRIC-seq) | H | HeLa | 1 | [45] |
| Akimitsu2 | BrU+4sU (Dyrec-seq) | H | HeLa | 1 | [46] |
| Bazzini | 4sU (SLAM-seq), ActD | H | K562, RPE, HeLa, HEK293 | 4 | [29] |
| Cramer | 4sU (TT-seq) | H | K562 | 1 | [47] |
| Cramer2 | 4sU (TT-seq) | H | K562 | 1 | [43] |
| Darnell | ActD | H | HepG2 | 1 | [38] |
| Dieterich | 4sU | H | HEK293, MCF-7 | 4 | [48] |
| Gejman | 4sU | H | LCLs (7 lines) | 15 | [44] |
| He | ActD | H | HeLa | 2 | [36] |
| Jaffrey | ActD | H | HEK293 | 1 | [42] |
| Marks | ActD | H | HeLa | 1 | [49] |
| Mortazavi | 4sU (Long-TUC-seq) | H | GM12878 | 1 | [50] |
| Oberdoerffer | BrU (BRIC-seq) | H | HeLa | 1 | [51] |
| Rinn | ActD | H | K562, H1-ESC | 6 | [52] |
| Rissland | 4sU | H | HEK293 | 4 | [27] |
| Shendure | 4sU | H | A549 | 1 | [53] |
| Rissland2 | 4sU, ActD, α-Amanitin | H, M | HEK293, 3T3 | 6, 2 | [41] |
| Simon | 4sU (TimeLapse-seq) | H, M | K562, MEF | 2, 2 | [54] |
| Zimmer | ActD | H, M | B cell, 3T3 | 1, 1 | [55] |
| Ameres | 4sU (SLAM-seq) | M | mESC | 1 | [56] |
| Bartel | ActD | M | 3T3 | 1 | [7] |
| Bartel2 | 5EU | M | 3T3 | 2 | [57] |
| Darnell | ActD | M | mESC | 1 | [58] |
| Dolken | 4sU (SLAM-seq) | M | M2-10B4 (bone marrow) | 1 | [59] |
| Hanna | ActD | M | mESC, mEB | 2 | [60] |
| Hu | ActD | M | mESC (E14Tg2a) | 1 | [61] |
| Ko | ActD | M | mESC | 4 | [6] |
| Koszinowski | 4sU | M | 3T3 | 1 | [62] |
| Mattick | ActD | M | Neuro-2a | 1 | [45] |
| Regev | 4sU | M | Dendritic cells | 1 | [63] |
| Regev2 | 4sU | M | Dendritic cells | 1 | [64] |
| Selbach | 4sU | M | 3T3 | 2 | [65] |
| Wilusz | ActD | M | C2C12 | 3 | [66] |

In order to evaluate the consistency among studies, we computed the pairwise Spearman correlations of all datasets, using the subset of common genes measured by each pair of studies. Most pairs of human datasets exhibited strong (i.e., ≥0.6) correlations (**Fig. 1**). However, the datasets segregated largely into two clusters; the first encompassed a diverse array of studies, cell types, and procedures, and the second encompassed datasets from only five studies [27,41–44] which appeared to be outliers in that they exhibited poor (i.e., ≤0.5) correlation to most studies other than to those from their own batch (**Fig. 1**). Most subclusters did not cleanly segregate by cell type or experimental method; rather, they tended to cluster closely by their study or laboratory of origin. This indicated the likely presence of batch effects which masked the cell-type specific signal captured by the data. Datasets from the mouse exhibited similar patterns of clustering, except that only two studies [60, 64] appeared to be the greatest outliers due to their poor correlation to most other studies (**Additional file 1: Fig. S2b**).

**Figure 1.**
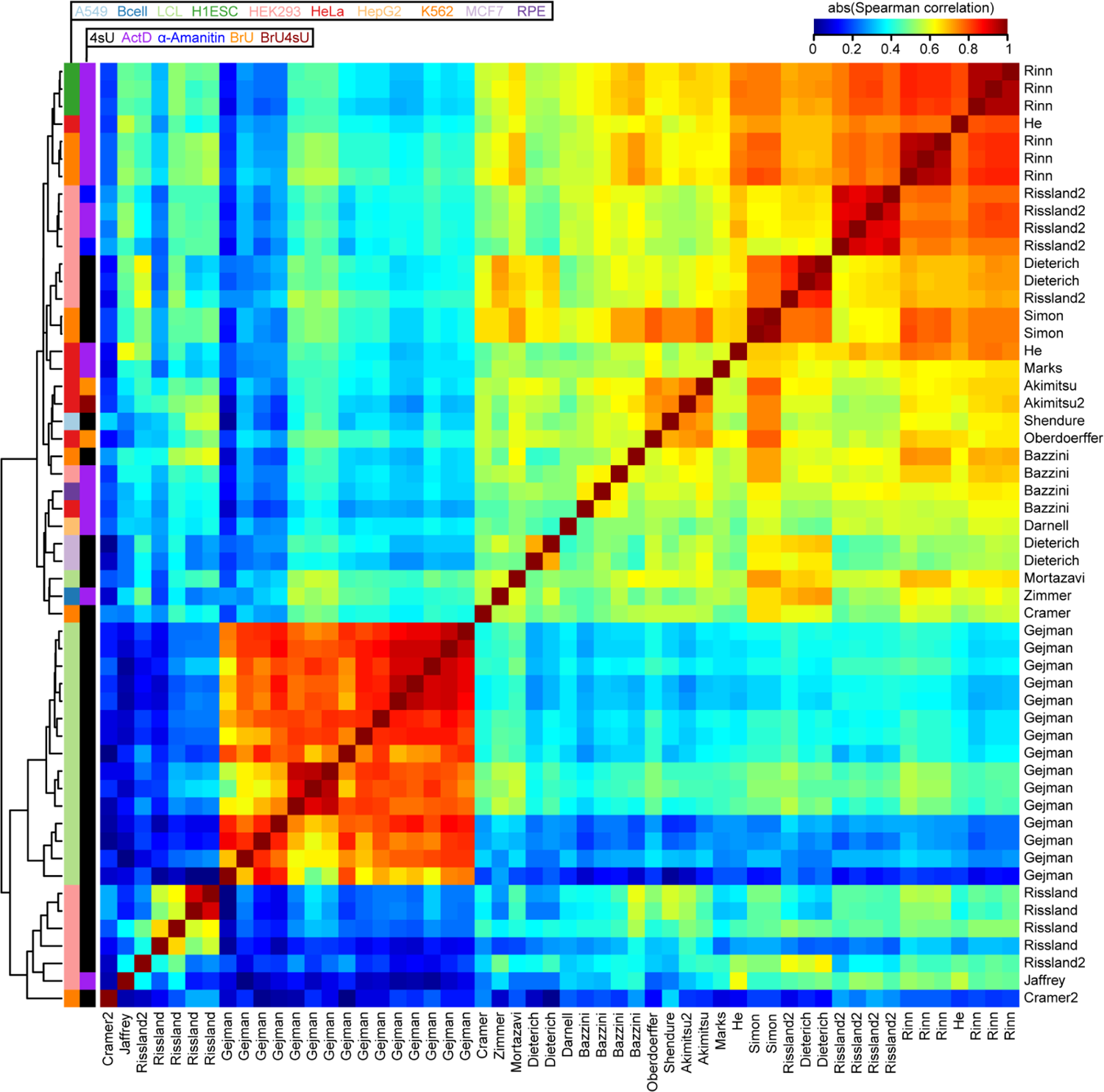
Comparison of half-lives in a compendium of human datasets. Heatmap of the absolute value of the Spearman correlations measured between half-lives derived from each pair of 54 human samples. Absolute values were used to accommodate five samples from four studies [29,38,46,49] whose data were deposited as degradation rates rather than half-lives. Samples are clustered using hierarchical clustering according to the indicated dendrogram. Rows are labeled by the study of origin (**Table 1**) and colored by the cell type of origin and measurement approach.

### Comparison of methodological bias and cell-type specificity captured in the mammalian half-life compendium

Given the wide disparity in reported genes (**Additional file 1: Fig. S1** and **S2a**) and potential existence of outliers in some samples, we pre-processed our *gene* x *sample* human and mouse half-life matrices to improve our ability to evaluate sample relatedness and examine possible sources of measurement bias. We standardized the samples in each matrix, used iterative PCA to impute missing gene measurements, and performed quantile-normalization to align the samples into similar distributions. In total, we recovered 13,921 human genes and 14,463 mouse genes in our matrices (**Additional file 3: Dataset S3**). Finally, we performed PCA to evaluate the relatedness among the 54 human samples. As observed previously (**Fig. 1**), PC1 identified samples from a single study [44] to be the greatest outliers relative to all other studies (**Additional file 1: Fig. S3**). We therefore found it parsimonious to assume that measurements from this study were severely biased, and henceforth excluded samples from this study from further analysis.

Reperforming the PCA analysis on the remaining 39 human samples revealed weak to non-existent clustering by cell type, and stronger clustering by measurement method (**Fig. 2a**). Specifically, PC2 seemed to segregate samples derived from pulse labeling experiments (i.e., those using 4sU, BrU, and BrU4sU) with those derived from transcriptional shutoff experiments (i.e., those using ActD and α-Amanitin). After averaging the PC2 values among replicates, we observed a statistically significant difference between the PC2 distributions for these two methodological classes (**Fig. 2b**), revealing a fundamental inconsistency in these techniques.

**Figure 2.**
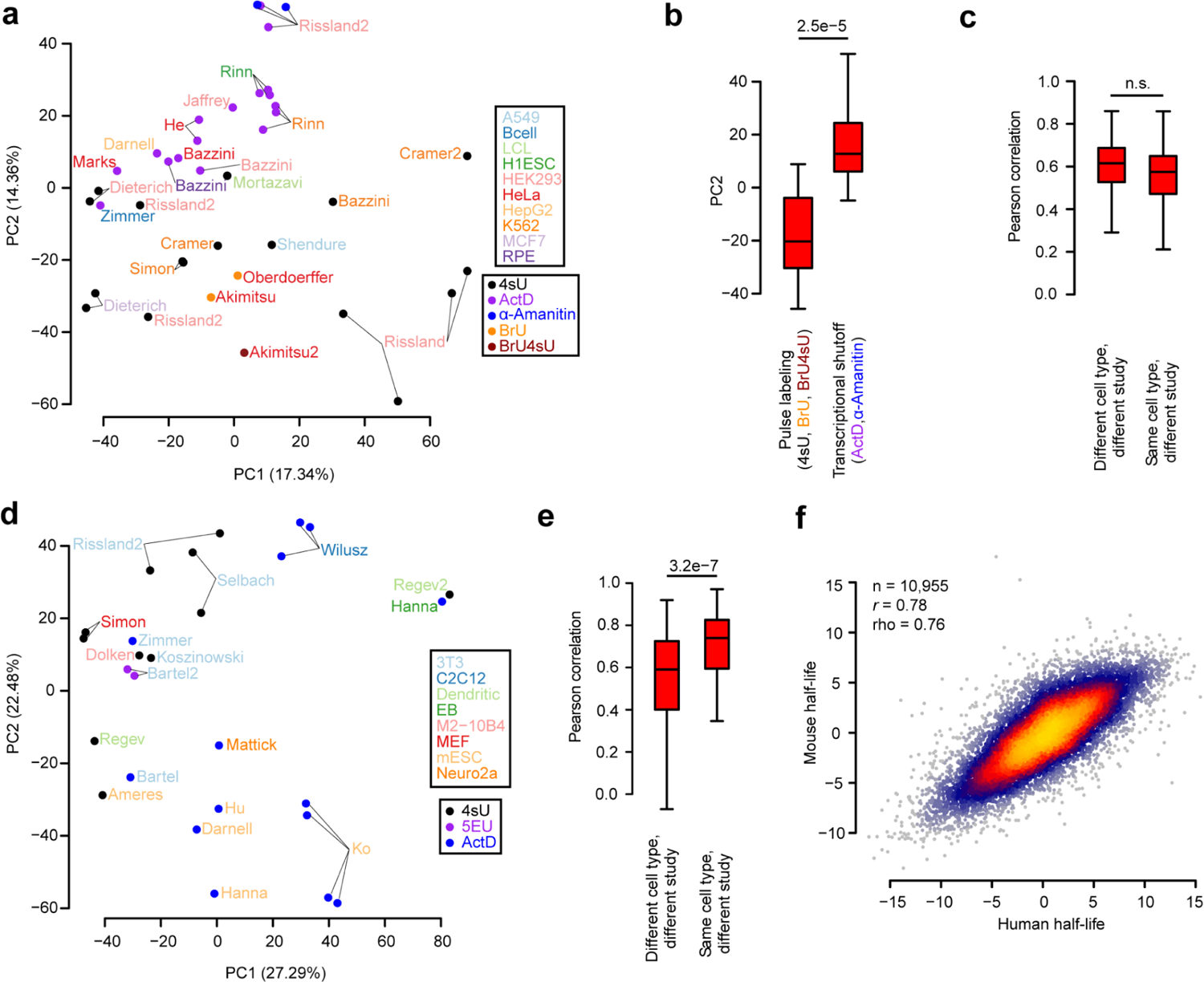
Assessment of measurement bias and cell-type specificity present in half-life data. **a)** PCA of all human samples except those from an outlier study [44], with sample names colored according to cell type and corresponding data point colored according to measurement approach. Axes are labeled according to the percentage of variance among samples explained by the first two PCs. See also **Additional file 1: Fig. S3** for the same analysis using all samples. **b)** Boxplot of sample distributions along PC2, partitioned according to the measurement method (i.e., pulse labeling or transcriptional shutoff). Replicates for the same study were first averaged according to their PC2 value prior to assessing differences between the methods, with statistical differences between distributions evaluated using a two-sided Wilcoxon rank-sum test. **c)** Evaluation of the Pearson correlations between pairs of half-life samples. Considered in this plot were the subset of pairs of two different studies that interrogated half-lives in either the same cell type or different cell types. Statistical differences between the distributions were evaluated using a one-sided Wilcoxon rank-sum test to assess whether correlations from the same cell type exceeded those from a different cell type. **d-e)** These panels are the same as those in (a) and (c), respectively, except compare mouse samples. **f)** Comparison of consensus, cell-type agnostic (i.e., methodology and cell-type independent) measurements of human and mouse half-lives among one-to-one orthologous genes. Half-lives for each species were computed as PC1 of the respective *gene* x *sample* matrix. Also indicated are the Pearson (r) and Spearman (rho) correlation values as well as sample size (n) of genes considered. Shown in all boxplots is the median value (bar), 25th and 75th percentiles (box), and 1.5 times the interquartile range (whiskers).

To ascertain whether the human data captured cell-type-specific half-lives, we evaluated all pairwise Pearson correlations between half-lives from samples derived from different studies (in order to minimize the influence of inflated correlations among replicates of the same study), but assessed on either the same cell type or different cell types. We hypothesized that we should see stronger correlations among samples evaluating the same cell type; however, we did not observe statistical support for this hypothesis (**Fig. 2c**), indicating that there was no detectable cell-type-specific signal for half-lives in human cells. Nearly identical results were achieved using Spearman correlations instead of Pearson correlations (data not shown). Given the known tissue-specific roles of miRNAs [67] and other post-transcriptional RBP regulators, we find it more plausible that technical noise and methodological bias obscure the cell-type specificity of half-life measurements relative to the possibility that none exists biologically.

Next, we again used PCA to evaluate the relatedness among the 27 mouse samples. In the mouse, there appeared to be stronger clustering by cell type and measurement methodology along both PC1 and PC2 (**Fig. 2d**). Indeed, Pearson correlations between pairs of half-life measurements from the same cell type and different study exceeded those from different cell types and different studies (**Fig. 2e**), suggesting the detection of a cell-type specific signature. Although this observation is encouraging, the limited sample size, coupled with the confounded nature of measurement methodology and cell type, make it ultimately impossible to decompose the relative influence of methodological bias and cell-type specificity in the measurement of mouse half-lives. In the future, a controlled study measuring half-lives in a number of different cell types with multiple experimental approaches would help to more accurately assess this.

Given our lack of success in identifying clear cell-type-specific measurements of half-life in the human and mouse samples, we instead focused on deriving cell-type-agnostic (i.e., universal) measures of half-life which integrate information across all methodologies and cell types. Towards this end, we computed the first PC in each of our *gene* x *sample* matrices to derive consensus measurements of half-life for each gene from each species. The PC1s explained 62.4% and 62.6% of variance among genes in each species, respectively, while PC2s explained merely 6.6% and 9.8% of variance, so were henceforth not used. Given that this procedure was performed independently in each species, we would expect that any artifacts induced by the procedure would corrupt the evolutionary relationship of measurements between species. However, we instead observed a strong concordance between our consensus half-life measurements among one-to-one orthologs between the two species (**Fig. 2f**). Our interspecies Pearson correlation of 0.78 greatly exceeds that of 0.61 reported previously in the literature [6]. This indicates that mRNA half-life has been more strongly conserved than previously appreciated between mammalian species separated by ∼75 million years of evolutionary time. Moreover, this finding also demonstrates our ability to recover a highly precise measurement of half-life that successfully ameliorates the impact of technical noise and methodological bias.

### A genetic model of mRNA half-life

Given our consensus measurements of mRNA half-life, we sought to determine whether it was possible to improve the predictability of half-life from mRNA sequence. Towards this end, we engineered groups of features (**Table 2**) with the goal of deriving a machine learning (ML)-based regression model to automatically select the subset of pertinent features. Our sequence-derived feature groups consisted of: i) basic mRNA features such as the length and G/C content of different functional regions and ORF exon junction density [2,6,7]; ii) *k*-mer frequencies of length 1-7 in the 5′ UTR, ORF, or 3′ UTR; iii) codon frequencies; iv) predicted repression scores of mammalian-conserved miRNA families [24]; and v) predicted binding of various RBPs to the mRNA sequence by SeqWeaver [68] and DeepRiPE [69]. RBP binding was predicted separately for the 5′ UTR, ORF, and 3′ UTR because it has previously been observed from cross-linking and immunoprecipitation sequencing (CLIP-seq) that RBPs have a tendency to bind in specific functional regions [70].

Next, we sought to determine which of these sequence-derived groups of features would be useful for predicting half-life. Because of the hierarchical association of each feature to a group (**Table 2**), we thus evaluated a series of nested models which iteratively considered additional groups of features. Each feature set was fed into a lasso regression model, and the relative model performance was compared between a simpler and more complex group on different folds of held-out data using a 10-fold cross-validation (CV) strategy. This helped to establish whether the inclusion of additional groups was justified. Evaluating the model series on our human half-life data, we observed that: i) 5′-UTR *k*-mers did not improve the model, ii) ORF *k*-mers did not improve the model beyond the simpler consideration of codons, iii) both 3′-UTR *k*-mers and predicted miRNA repression scores improved the model, and iv) SeqWeaver predictions of RBP binding improved the model more than those from DeepRiPE (**Fig. 3a**). Our final model that optimally balanced the tradeoff between complexity and performance was the “BC3MS” model, which considered basic mRNA features, codon frequencies, 3′-UTR *k*-mers, miRNA repression scores, and SeqWeaver predictions.

**Figure 3.**
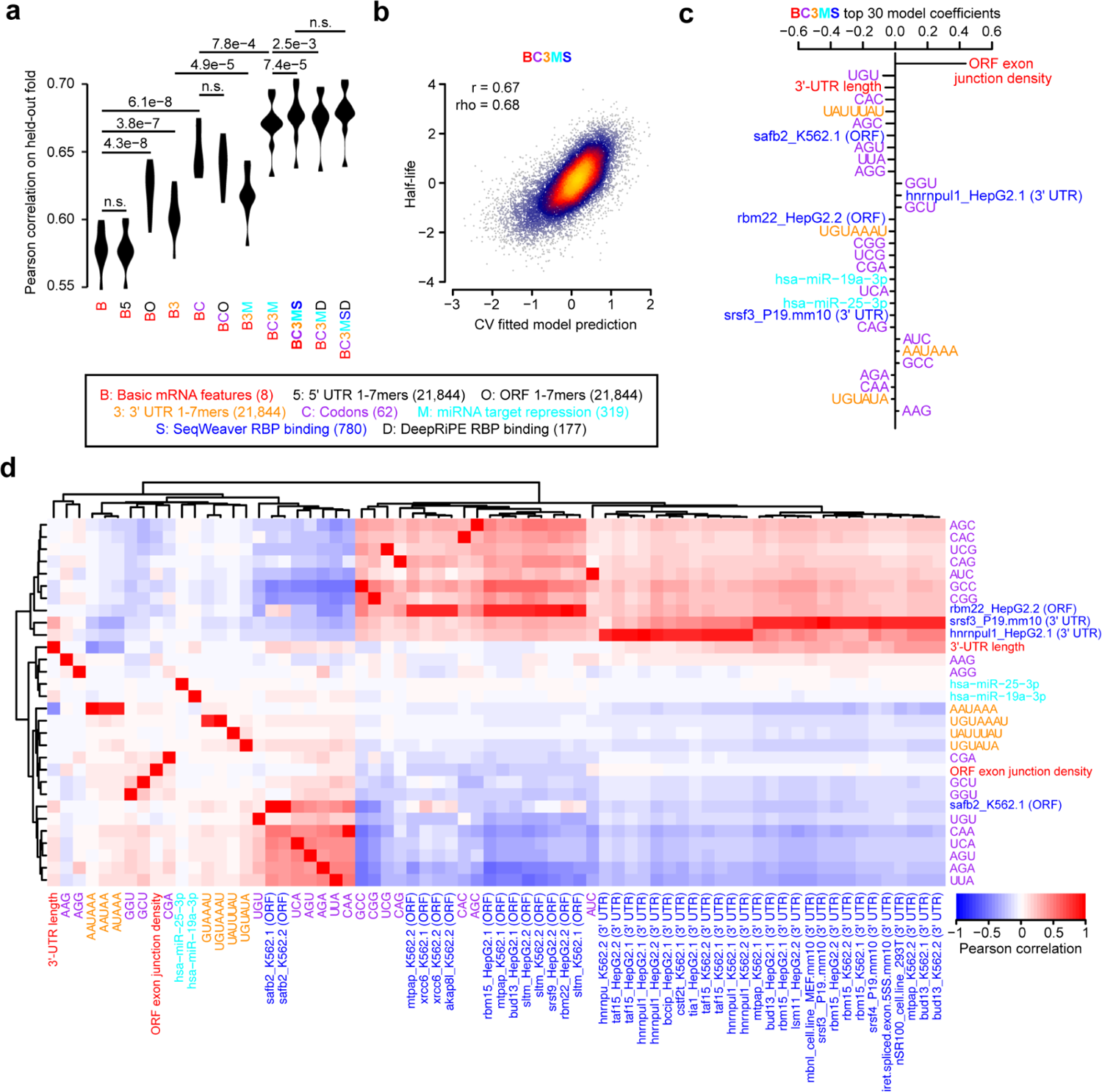
Prediction of human half-lives using sequence-encoded features. **a)** Performance of trained lasso regression models on each of 10 held-out folds of data. Compared is the relative performance between pairs of nested models which iteratively consider greater numbers of features. Each model is described by a code indicating the features considered. A description of the code is provided in the key, along with the corresponding number of features considered listed in parentheses. An improvement in a more complex model relative to a simpler model was evaluated with a one-sided, paired t-test, adjusted with a Bonferroni correction to account for the total number of hypothesis tests. Features which were ultimately determined to contribute to performance improvement are colored, or are left black if they did not improve the model. **b)** Shown are the final predictions for the optimal model (i.e., BC3MS) after concatenating the observations for all 10 folds of held-out data. Also indicated are the Pearson (r) and Spearman (rho) correlation values. **c)** The top 30 ranked model coefficients corresponding to the BC3MS model, trained on the full dataset. Features are colored according to the same key as that in panel (a). **d)** Pearson correlation matrix between the union of all top 30 features from (c), shown as rows, and other features sharing a Pearson correlation either ≤ –0.8 or ≥ 0.8, shown as columns. Feature names are colored according to the origin of the feature as shown in the same key as panel (a). Hierarchical clustering was used to group features exhibiting similar correlation patterns.

**Table 2.** Summary of features considered in models trained to predict mRNA half-life, with a description of the features considered, feature type (*i.e.*, sequence or biochemical), data source, and number of features in the category. Sequence features were calculated identically for both human and mouse. Biochemical features were calculated only with respect to human data.

| Feature description | Feature type | Data source | # features |
| --- | --- | --- | --- |
| Basic mRNA features such as length and G/C content of 5' UTR, ORF, and 3' UTR; intron length; ORF exon junction density | Sequence-derived | Custom Perl scripts | 8 |
| Codon frequencies | Sequence-derived | Custom R scripts | 64 |
| 1- to 7-mer frequencies in the 5' UTR | Sequence-derived | Custom R scripts | 21,844 |
| 1- to 7-mer frequencies in the ORF | Sequence-derived | Custom R scripts | 21,844 |
| 1- to 7-mer frequencies in the 3' UTR | Sequence-derived | Custom R scripts | 21,844 |
| Predicted average binding score of human RBPs (in each of 5' UTR, ORF, and 3' UTR) | Sequence-derived | DeepRiPe predictions [69] | 177 |
| Predicted average binding score of human and mouse RBPs (in each of 5' UTR, ORF, and 3' UTR) | Sequence-derived | SeqWeaver predictions [68] | 780 |
| Number of CLIP peaks for various RBPs | Biochemical | ENCORI database [71] | 133 |
| Number of eCLIP peaks for various RBPs (K562, HepG2, and adrenal gland) | Biochemical | ENCORE database [72] | 225 |
| Number of PAR-CLIP peaks | Biochemical | [70] | 146 |
| Number of CLIP peaks for m6A pathway components (a6A, m6Am, YTHDF2, METTL3, METTL14, and WTAP) | Biochemical | [37,42,73–77] | 13 |
| RIP-seq of diverse RBPs and translational efficiency | Biochemical | [36,78–80] | 34 |

Concatenating the predictions across all folds of held-out data, we observed a correlation of 0.67 between the BC3MS model predictions and observed human half-lives (**Fig. 3b**). We retrained this model on the full dataset to assess which features contributed most significantly to half-life prediction. The top-30 ranked features of the model consisted primarily of basic mRNA properties, codon frequencies, and predicted miRNA/RBP binding sites (**Fig. 3c**). Consistent with previous work [2,6,7], ORF exon junction density was the dominant feature. Moreover, the signs of coefficients associated with codon frequencies are consistent with previously published human Codon Stability Coefficients (CSCs) [26,27,29,30], which were computed based upon an isolated Pearson or Spearman correlation between the codon frequencies and half-lives [28]. CSCs have been previously established as a quantitative metric that captures the association between a codon and mRNA stability [26–30]. Our model contextualized relative codon influence with respect to other sequence features in a multiple linear regression framework, and re-ranked their utility in the prediction task.

Next, despite considering over 20,000 possible 3′-UTR *k*-mers, the model automatically discovered highly conserved regulatory motifs such as UAUUUAU, the core AU-rich element (ARE) [16, 34]; UGUAAAU and its variant UGUAUA, Pumilio binding elements [33]; and AAUAAA, the cleavage and polyadenylation motif involved in alternative polyadenylation [81, 82]. The binding of four RBPs emerged as the most useful features, including that of SAFB2 and RBM22 in the ORF as well as HNRNPUL1 and SRSF3 in the 3′-UTR. Finally, consistent with previous work showing that miRNAs weakly impact half-life [7], the model predicted light repressive roles for two miRNAs, miR-19a and miR-25.

Although we find these factors to be promising candidates, we caution that the interpretation of feature selection and coefficient-based ranking is inherently limited by the substantial degree of multicollinearity among features. To guard against the possibility of lasso regression selecting one feature over another through its spuriously higher correlation to half-life, we examined the full set of features strongly correlated to the top 30 selected features (**Fig. 3d**). While the majority of features were not strongly correlated to other features, we indeed found that SeqWeaver predicted a similar degree of binding among numerous alternative factors such as MTPAP, XRCC6, AKAP8L, RBM15, BUD13, SLTM, SRSF9, HNRNPU, TAF15, BCCIP, CSTF2T, TIA1, LSM11, MBNL, and SRSF4. We consider this full set of RBPs, in conjunction with our earlier set, as candidate post-transcriptional regulators for future experimental investigation.

We repeated the same analyses independently for the mouse, with the goal of devising a genetic model to predict mouse half-lives. While most findings were highly similar between the two species, key differences include: i) DeepRiPE features significantly boosted performance, leading to the optimality of the “BC3MSD” model (**Additional file 1: Fig. S4a**); ii) the model achieved a slightly lower Pearson correlation of 0.61 (**Additional file 1: Fig. S4b**); iii) the feature ranking varied, with the inclusion of FMR1, SRSF4, HNRNPM, RBM15, IGF2BP3, KHSRP, MBNL, miR-27a, and FXR2 in the top 30 coefficients (**Additional file 1: Fig. S4c**); and additional factors correlated with the top 30 features, such as HNRNPM, SFPQ, SRSF3, TRA2A, SRSF1, EIF3D, DDX6, XRCC6, and SRSF7 (**Additional file 1: Fig. S4d**).

Finally, we evaluated the ability of the models trained in each species to generalize to the opposite species. Our interspecies comparisons revealed that models tested in the opposite species performed competitively, albeit slightly worse, than models trained within the same species, for both the human and mouse (**Additional file 1: Fig. S4e**). This indicates that the vast majority of the learned half-life-associated features have strong predictive value more generally across the mammalian phylogeny.

### A biochemical model of mRNA half-life

Given the recent trove of biochemical data evaluating RBP binding, we attempted to identify experimentally supported features that are predictive of mRNA half-life. Towards this goal, we assembled an array of several large-scale datasets measuring RBP binding to develop an ML-based regression model analogous to our genetic model. Our biochemical feature groups consisted of: i) the number of CLIP peaks in an mRNA for various RBPs in the ENCORI database [71], ii) the number of eCLIP peaks for various RBPs in the ENCORE database [72], iii) the number of PAR-CLIP peaks for various RBPs [70], iv) the number of CLIP peaks for m6A pathway components [37,42,73–77], v) and RIP-seq of diverse RBPs and translational efficiency [36,78–80] (**Table 2**). Implementing the same strategy as before to compare nested models with 10-fold CV, we observed that: i) PAR-CLIP data did not improve the model, ii) RIP-seq data did not improve the model after jointly considering eCLIP and m6A CLIP data, and iii) the remaining datasets all improved the model, with the greatest benefit emerging from considering the ENCORI database (**Fig. 4a**). “BEeM” was our best model, which considered basic mRNA features, ENCORI CLIP, eCLIP, and m6A CLIP data.

**Figure 4.**
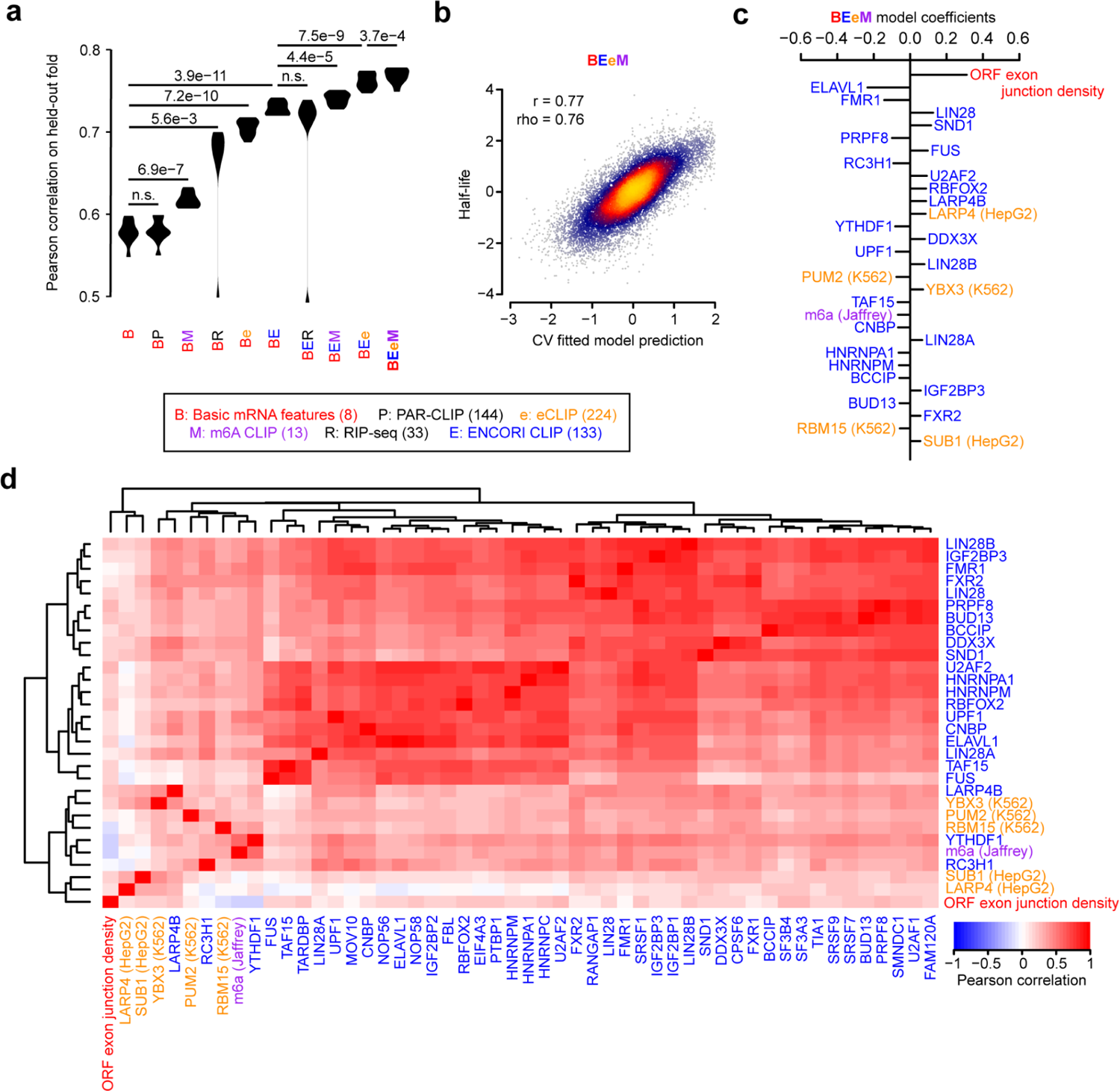
Prediction of human half-lives using biochemical features. This figure is organized in the same fashion as **Fig. 3**, except it evaluates features derived from biochemical experiments. All CLIP data is computed as the number of peaks on the full-length transcript, while RIP-seq is represented as a continuous measurement of the enrichment of RBP binding relative to a control IP.

We observed a global correlation of 0.77 between the BEeM model’s held-out predictions and observed human half-lives (**Fig. 4b**). The top-30 ranked features of the model (i.e., retrained on the full dataset) consisted primarily of the ORF exon junction density and dozens of RBPs (**Fig. 4c**). A number of RBPs consistently emerged between our genetic (**Fig. 3c,d** and **Additional file 1: Fig. S4c,d**) and biochemical models, including BUD13, BCCIP, HNRNPM, FMR1, PUM1 (i.e., a Pumilio-element binding protein), ELAVL1 (i.e., an ARE binding protein), TAF15, IGF2BP3, FXR2, and RBM15. Novel components that were not previously captured by the genetic model include LIN28A/B, SND1, PRPF8, FUS, RC3H1, U2AF2, RBFOX2, LARP4, YTHDF1 and its ligand m6A, DDX3X, UPF1, YBX3, CNBP, HNRNPA1, and SUB1 (**Fig. 4c**). Collectively, these factors were strongly correlated to others which serve as candidate regulators of mRNA stability (**Fig. 4d**).

Finally, we tested the possibility that a joint genetic and biochemical model might outperform either individually. A lasso regression model jointly trained upon BC3MS (i.e., genetic) and BEeM (i.e., biochemical) features modestly outperformed our BEeM model alone (**Additional file 1: Fig. S5a**), achieving a Pearson correlation of 0.78 (**Additional file 1: Fig. S5b**). The biochemical features were dominantly used in this model, although several codons were selected, along with the ARE 7-mer, which minimized the effect size of the coefficient attributable to ELAVL1 (**Additional file 1: Fig. S5c,d**).

### A deep-learning based genetic model of mRNA half-life

Having compared the performance of genetic and biochemical models, we sought to evaluate whether an alternative learning paradigm might be able to automatically decipher sequence-based rules that are more predictive of mRNA half-life. Towards this goal, we trained a hybrid convolutional and recurrent deep neural network architecture, called Saluki, which has the advantage of potentially uncovering nonlinear (e.g., cooperative) relationships among motifs and spatial principles with respect to motif positioning. The input to our model was a binary encoding of the mRNA sequence (i.e., up to a maximum of 12,288 nt), concatenated with binary variables for each nucleotide indicating the presence or absence of the first reading frame of a codon and splice site (**Fig. 5a**). This matrix was fed in to a neural network comprised of 64 1D convolutions (width 5), a max pooling layer (width 2), 6 additional blocks of these aforementioned layers, a recurrent layer consisting of a gated recurrent unit (GRU), and a densely connected layer (**Methods**, **Fig. 5a**, and **Additional file 1: Fig. S6a**). Given that mRNA sequences are variable in length, the 3′ end of each mRNA was padded with Ns after the transcriptional end site. To account for this property of the input, we left-justified the input matrix, and the GRU was oriented to pass information from the right to left such that the sequence was still most recently considered by the network (i.e., as opposed to the padded Ns) when integrating information from the full sequence to make a prediction. The parameters comprising this common “trunk” of the network were jointly trained on human and mouse data in alternating batches of species-specific data, with two “heads” to predict human and mouse half-lives, respectively [4, 5].

**Figure 5.**
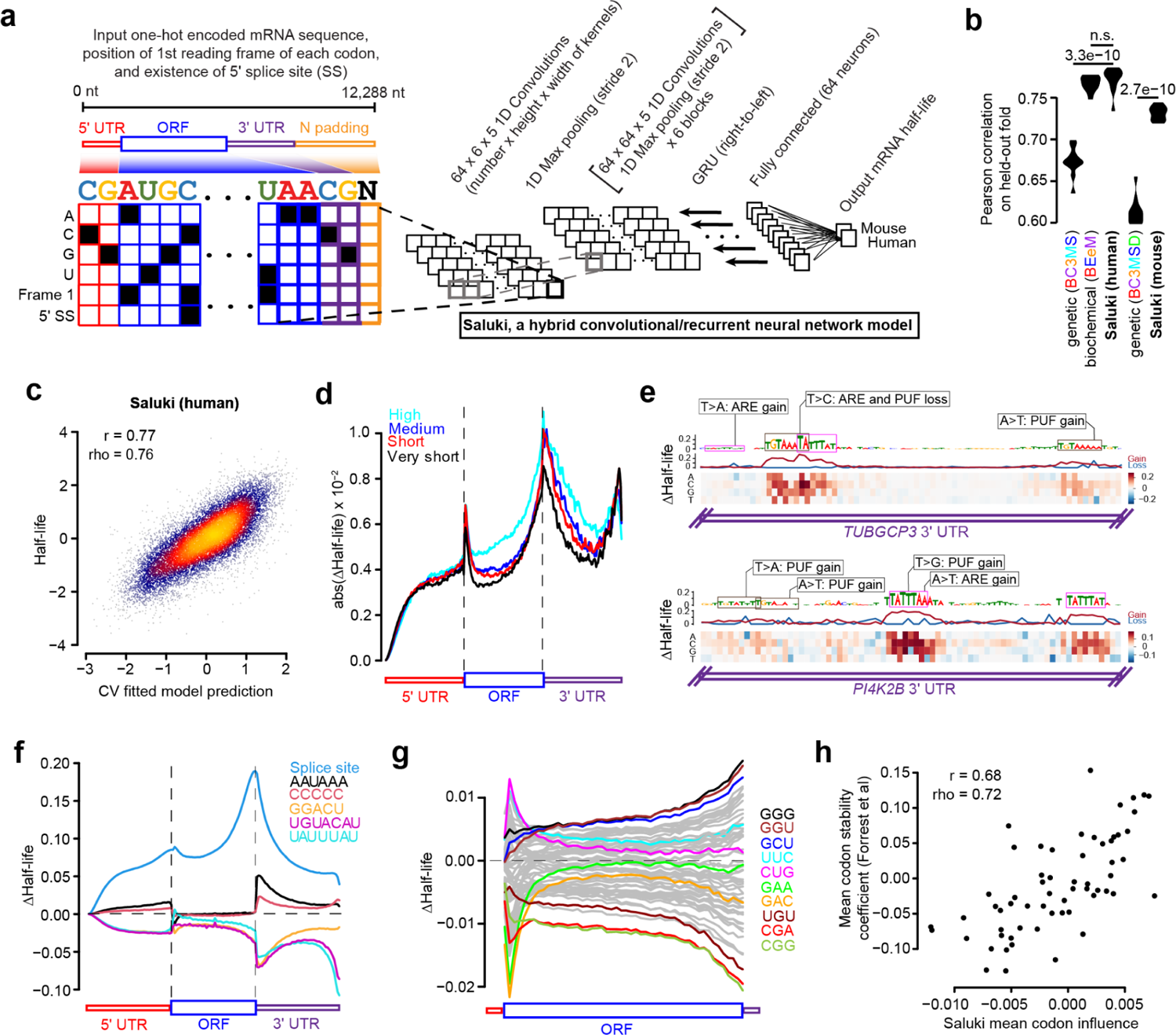
State-of-the-art prediction of half-lives and genetic variant functional effects using a sequence-based deep learning model. **a)** A hybrid convolutional/recurrent neural network architecture to predict half-life from an input of the RNA sequence, an encoding of the first frame of each codon, and 5′ splice site junction(s). The deep learning model, called Saluki, was jointly trained on mouse and human half-life data to predict species-specific half-lives. b) Performance of the trained Saluki models on each of 10 held-out folds of data, relative to the corresponding performances from our best genetic (i.e., “BC3MS” for human and “BC3MSD” for mouse, respectively) and biochemical (i.e., “BEeM”) lasso regression models. An improvement relative to another model was evaluated with a two-sided, paired t-test. c) Shown are the final predictions after concatenating the observations for all 10 folds of held-out data. Also indicated are the Pearson (r) and Spearman (rho) correlation values. d) Metagene plot of ISM scores across all mRNAs for percentiles along the 5′ UTR, ORF, and 3′ UTR. mRNAs were grouped into one of 4 bins according to their predicted half-lives. For the set of mRNAs within each bin, we plotted the average of the absolute value of the mean predicted effect size (i.e., of the three possible alternative mutations). e) ISM results of two 3′-UTR segments from *TUBGCP3* and *PI4K2B*. Partial matches to the AU-rich element (ARE, or “UAUUUAU”) and Pumilio/FBF (PUF, or “UGUAHAUA”) binding element consensus sequences are boxed. For each motif, single point mutations resulting in particularly severe or opposite phenotypes are shown alongside annotations reflecting the corresponding ARE and PUF consensus gain or loss events. f) Insertional analysis of motifs discovered by TF-MoDISco [83]. Each motif was inserted into one of 50 positional bins along the 5′ UTR, ORF, and 3′ UTR of each mRNA. Indicated is the average predicted change in half-life for each bin plotted along a metagene. g) This panel is the same as panel f), except it performs analysis of 61 codons (excluding the 3 stop codons) inserted into the first reading frame along the length of the ORF. Selected codons are colored, with the rest shown in gray. h) Scatter plot showing the relationship between the mean influence of each codon along the length of the ORF, as predicted by Saluki in panel g), and the mean codon stability coefficient over a set of cell types as observed previously [26]. Also indicated are the Pearson (r) and Spearman (rho) correlation values.

We performed an ablation analysis in which the model was evaluated after training it with only sequence as input; sequence and reading frame tracks; and sequence, reading frame, and splice site tracks. This analysis revealed that all forms of information were utilized by the network for Saluki to achieve optimal performance (**Additional file 1: Fig. S6b**). Training our Saluki model with the identical folds of data as our lasso regression models enabled us to directly compare their respective performances on each of the 10 folds of held-out data. Averaging the predictions using models derived from each of five independent training runs for each fold, Saluki’s performance exceeded that of the genetic lasso regression models for each species (i.e., “BC3MS” for human and “BC3MSD” for mouse), and it performed nearly identically with the human biochemical regression model (i.e., “BEeM”) (**Fig. 5b**). The final human and mouse models displayed correlations of 0.77 and 0.73, respectively (**Fig. 5c** and **Additional file 1: Fig. S6c**), suggesting that Saluki potentially learned novel principles of post-transcriptional gene regulation which our simpler linear models were unable to capture.

In the hopes of revealing such principles, we tested the predictive behavior of Saluki in different contexts. First, we performed an *in silico* mutagenesis (ISM) for every human mRNA in our dataset, mutating each reference nucleotide of an mRNA into every alternative allele to evaluate the predicted change in half-life [1,4,5,84]. We generated a metagene plot using these scores, evaluating the absolute value of the mean effect size (i.e., of the three possible alternative mutations) in percentiles along the length of each functional region (**Fig. 5d**). This analysis revealed that the model predicts a relatively modest influence for the 5′ UTR relative to the ORF and 3′ UTR. Moreover, the termini of functional regions, such as the 3′ terminus of the 5′ UTR as well as the 5′ and 3′ termini of the ORF and 3′ UTR, were revealed to harbor the most informative positions along an mRNA that contribute to half-life (**Fig. 5d**).

We further interrogated our ISM scores to identify the most pertinent motifs associated with changes in half-life using TF-MoDISco [83]. The poly-A binding element (“AAAAAAA”), ARE (“UAUUUAU”), PUF element (“UGUAHAUA”), putative ELAVL1/2 element (“UUUAU”), polyadenylation element (“AAUAAA”), and m6A motif (“GGACU”) were enriched as significant motifs in the 3′ UTR. Of these, the m6A motif and PUF element were also enriched in the ORF and 5′ UTR, and the putative ELAVL1/2 and polyadenylation elements were enriched in the 5′ UTR (**Additional file 1: Fig. S6d**). Visualizing ISM scores for two random examples, the 3′-UTR segments from *TUBGCP3* and *PI4K2B*, illustrates several learned properties involving these motifs (**Fig. 5e**). Broadly speaking, a mutation ablating a core ARE or PUF element led to a predicted increase in mRNA half-life. Conversely, several point mutations led to a gain of a motif, leading to a predicted decrease in half-life. While a subset of these mutations generated novel motifs, others changed an existing motif closer to its consensus sequence. In some cases, a mutation caused a dual loss of overlapping PUF/ARE motifs, leading to a more severe predicted change in half-life relative to the loss of each individual motif (**Fig. 5e**).

Next, we evaluated how the model would behave if we inserted either a splice site or one of our five enriched motifs along the full length of each mRNA. We performed this insertional analysis for most human mRNAs in our dataset, and then averaged the result according to the spatial bin of the insertion along each functional region. Consistent with their known roles, we observed that insertion of an m6A site, ARE, or PUF element reduced mRNA stability, with the greatest effect size arising from a 3′-UTR insertion (**Fig. 5f**). In contrast, insertion of a splice site, polyadenylation element, or C-rich element led to enhanced mRNA stability. Consistent with the lasso regression model, the presence of a splice site led to at least a four-fold enhancement in half-life relative to alternative motifs. The most novel property captured in the deep learning model—yet absent from our lasso model—is the strong dependence between the spatial coordinate of the motif along the mRNA and its predicted impact on mRNA half-life, both across and within functional regions (**Fig. 5f**). For instance, ARE and PUF sites are predicted to most strongly repress mRNA stability if they occur at the 5′ or 3′ terminus of the 3′ UTR, reminiscent of a well-known property of microRNA-mediated repression [24, 85]. In contrast, the m6A element is predicted to most destabilize mRNA immediately after the stop codon, mirroring the known relationship between m6A deposition and mRNA stability [58].

Given the mechanistic link between codon usage and mRNA half-life [8,26–30], we sought to ascertain whether Saluki has also learned this property. We therefore reiterated our insertional analysis, this time using 61 codons (excluding the 3 stop codons) inserted into the first reading frame along the length of each ORF. As before, the model attributed substantially different effect sizes to codons depending upon their position along an ORF, with the greatest predicted effects occurring close to the start and stop codons (**Fig. 5g**). Although most codons had the greatest predicted effect when inserted closer to the stop codon, several codons such as “CUG”, “UUC”, “GAA”, and “GAC” were predicted to have a modest effect when inserted into most regions except near the start codon (**Fig. 5g**). We sought to compare our predicted codon effects to existing measures of codon influence, such as the aforementioned CSCs [26–30]. We therefore investigated the relationship between the mean codon influence across an ORF, as predicted by Saluki, relative to the mean CSC across numerous cell types as quantified previously [26]. Reassuringly, there was a strong relationship between the two metrics (Pearson correlation = 0.68, **Fig. 5g**), suggesting that Saluki successfully captures the influence of codon usage.

### Prediction of 3′-UTR regulatory function and genetic variant effects

Given Saluki’s strong performance in predicting endogenous half-lives, we sought to evaluate its ability to predict the effect of mRNA sequence and genetic variants therein on mRNA stability in a more controlled context. Massively parallel reporter assays (MPRAs) provide an ideal setting to test causal relationships, because they can directly assess how specific sequences influence reporter expression. Several studies have deployed MPRAs to test the functional impact of thousands of 3′-UTR fragments and genetic variants therein on mRNA stability [86–88]. We performed *in silico* versions of these MPRA experiments to evaluate their consistency with *in vivo* experiments.

The first study performed three types of MPRA experiments: i) evaluating the impact of mutation on 8-nt intervals tiling the *CXCL2* 3′ UTR, ii) performing a saturation mutagenesis of a specific region within the same 3′ UTR to measure variant effects, and iii) testing the effect of 3,000 highly conserved 3′-UTR segments on RNA stability [86]. To compare our predictions to the first of the three MPRA experiments, we compared our pre-computed ISM scores for the *CXCL2* 3′ UTR to those observed in the experiment (**Fig. 6a**). We observed a general qualitative agreement between model predictions and experiment in the 3′-UTR regions showing high activity, which typically overlapped conserved and ARE-containing regions (**Fig. 6a**). However, there were several novel elements detected by the assay which were conserved but not predicted by the model, indicating several false negatives among the predictions. Next, we evaluated how well our predictions agree with saturation mutagenesis data from the tested *CXCL2* 3′-UTR region. To more closely simulate this experiment *in silico*, we integrated each of the measured oligonucleotide fragments into the 3′ UTR of eGFP within the corresponding BTV vector tested in the experiment [86]. Then, for each mutation, we calculated the predicted variant effect as the divergence of predicted half-lives between the reference and alternative sequence. We observed a strong agreement between the observed and predicted variant effects, in which both methods highlighted a strong effect for a set of upstream AREs and a weaker effect for a downstream ARE (**Fig. 6b**). Collectively, the variant effect predictions agreed with those observed with a Spearman correlation of 0.69 (**Fig. 6c**). Finally, we assessed how well we could predict the observations of an MPRA testing the effect of 3,000 highly conserved 3′-UTR segments on RNA stability. We achieved a Spearman correlation of 0.63 between our model predictions and experiment (**Fig. 6d**), suggesting that our model is able to integrate the causal relationship between RNA sequence and RNA stability.

**Figure 6.**
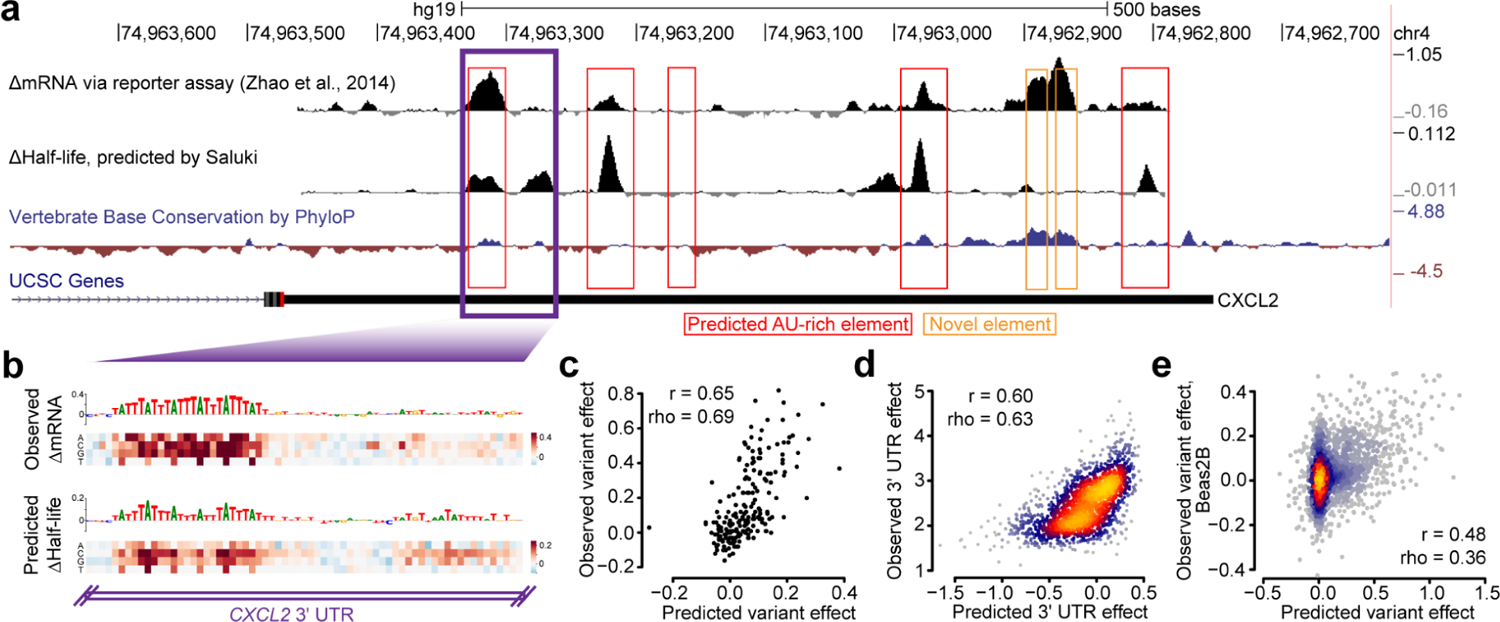
Concordance of Saluki predictions and functional data from massively parallel reporter assays. **a)** Effect of mutation on RNA stability, as measured by an MPRA [86], for tiles along the *CXCL2* 3′ UTR separated by 8-nt intervals. Also shown are variant effect predictions from Saluki (smoothed along a local 8-nt window) for the same region, and vertebrate base conservation as measured by PhyloP [89]. Predicted AREs are boxed in red, and novel elements detected by the MPRA are boxed in orange. **b)** Saturation mutagenesis of a segment of the *CXCL2* 3′ UTR, boxed in purple in part (a). Shown are the observed variant effects (top) and Saluki’s predicted variant effects (bottom). The reference sequence is shown for each, in which the nucleotide height is scaled according to the mean observed or predicted effect for that position. **c)** Scatter plot of the observed and predicted variant effects shown in panel (b). **d)** Scatter plot of the observed and predicted 3′-UTR effects for each of 3,000 conserved 3′ UTRs profiled by fastUTR [86]. **e)** Scatter plot of the observed and predicted variant effects, as measured in Beas2B cells [87]. Also indicated are the Pearson (r) and Spearman (rho) correlation values for panels (c-e).

To further evaluate the generality of our model, we turned to other MPRAs testing both 3′-UTR and variant effects in Jurkat and Beas2B cells; the elements tested were heavily enriched in ARE-containing regions [87]. We observed Spearman correlations of 0.42 and 0.49 between our model predictions and observed 3′-UTR effects in Jurkat and Beas2B cells, respectively (**Additional file 1: Fig. S7a,b**). These predictions were similar to the observed 3′-UTR effects between both cell types (Spearman correlation = 0.52, **Additional file 1: Fig. S7c**), suggesting that the model was about as predictive as the same MPRA experiment performed in different cell types. Similarly, when evaluating variant effects in both cell types, we achieved Spearman correlations of 0.31 and 0.36 between our model predictions and observed variant effects in Jurkat and Beas2B cells, respectively (**Fig. 6e** and **Additional file 1: Fig. S7d,e**). Yet again, this predictive performance was on par with the agreement of observed variant effects between the two cell types (Spearman correlation = 0.26, **Additional file 1: Fig. S7f**).

As a final test for the model, we evaluated its agreement with MPRAs that measured the functional effect of 12,173 3′ UTRs in six cell types [88]. While the observed values agreed well between any pair of cell types (Spearman correlations from 0.60 to 0.83), our predictions agreed a bit more modestly (Spearman correlations from 0.26 to 0.50, **Additional file 1: Fig. S7f**), suggesting that they partially captured the causal factors linking RNA sequence to stability in this dataset.

## DISCUSSION

Since the emergence of the first transcriptome-wide measurements, there have been numerous efforts to understand the sequence-encoded determinants of mRNA half-life [6–8,38–40,90,91].

Despite the diversity of modeling approaches developed to predict half-life, the quality of the predictive engines has been severely limited by the quality of the measurement itself. It has long been recognized that different methodologies suffer from biased measurement [16, 17], leading to an inconsistent portrait of transcriptome-wide half-lives. Indeed, nearly three decades ago, it was theorized that considering an ensemble of different methods would ultimately lead to a more precise definition of half-life [18]. In this study, we aimed to establish a more precise “ground truth” of mammalian mRNA half-lives, empowered by the systematic collection and subsequent meta-analysis of a large cohort of mammalian half-life datasets. As anticipated, this exercise revealed inconsistencies among data derived from different research groups and experimental protocols (e.g., pulse-chase vs transcriptional shutoff). We therefore derived consensus measurements of mRNA half-life which were less susceptible to technical noise and methodological bias.

Importantly, these bias-adjusted measurements led to several favorable outcomes. First, there was an improved interspecies agreement between the half-lives of orthologous human and mouse genes, indicating a greater conservation of this molecular trait than previously appreciated. Next, there was a substantial improvement in quantitative efforts to predict half-life from genetic and biochemical features. Finally, the relative importance of the heterogeneous mechanisms influencing half-life could be better delineated. Our lasso regression and deep learning models captured most of the known determinants of half-life, integrating the influence of AREs, PUF elements, YTHDF1-binding elements, splice sites, microRNA binding sites, and codon composition. While our lasso models implicate novel roles for several additional RBPs, our deep learning models propose novel properties of gene regulation such as the position-dependent effects of codons, motifs, and splice sites.

Aside from its improved accuracy, one of the key advantages of our deep learning model relative to our lasso regression models is its ability to rapidly generate predictions given arbitrary sequences, obviating the need to compute hundreds to thousands of features for each mRNA. This property makes Saluki amenable to the rapid prediction of variant effects, enabling us to produce a global map of predicted changes in half-life for all possible mutations in the exons of protein-coding genes. Given the concordance between our *in silico* predictions as well as *in vivo* MPRA measurements of regulatory element and variant function, our work offers a promising initial foray into the problem of predicting the post-transcriptional consequences of genetic mutation in exonic sequences.

## CONCLUSIONS

In this study, we demonstrated our ability to predict mRNA half-life from sequence with an r^2^ of 0.59, which represents nearly a 50% performance improvement relative to existing models in mammals (r^2^=0.20 and r^2^=0.39) [6, 7]. Among the biggest reasons for this improvement is simply the derivation of a more precise measurement of mRNA half-life from a meta-analysis of transcriptome-wide half-life datasets. Given our encouraging results in the auxiliary tasks of sequence and variant effect prediction, as quantified by the consistency between model predictions and MPRA data, we foresee several practical applications for Saluki.

First, there is a critical need to bioengineer stable RNAs for vaccine development and gene therapy applications. Despite the recent success of mRNA vaccines targeting Covid-19, they are inherently unstable and prone to degradation due to the prevalence of intracellular and extracellular RNases [92]. Problems surrounding mRNA instability and inefficient protein expression have been partially overcome through the incorporation of stabilizing mRNA structures [15], a 5’-cap, modified nucleosides, and codon optimality rules [92]. In the context of adeno-associated virus (AAV)-based gene therapy vectors, codon optimality rules have also been engineered to encourage greater mRNA stabilities and higher translation rates for several therapeutic proteins [93]. Nevertheless, the global sequence has not yet been jointly optimized for the mRNA to exhibit a longer half-life. Deep learning models have been successfully coupled to generative models, such as generative adversarial networks, to engineer nucleic acids to obey certain properties. This is exemplified by the design of 5′ UTRs to improve translation rate [94], 3′ UTRs to enhance cleavage and polyadenylation [95, 96], and coding sequences to evolve antimicrobial peptides [97]. Similarly, we envision Saluki’s use as an external function analyzer, or “oracle”, to engineer mRNAs with desired half-life properties.

Given the widespread roles of mRNA decay in the genetics of human health and disease [12,88,98], Saluki is also poised to provide insight into whether a genetic variant functions through post-transcriptional gene regulatory mechanisms. Towards this goal, it could help to fine-map causal eQTLs through the integration of its score into supervised learning frameworks [13]. Such scores could also be integrated into tools intended to identify pathogenic non-coding variants in the human genome [14] to further unravel the genetic basis of disease.

## METHODS

### mRNA half-life data collection and pre-processing

We manually collected the processed half-life values from all of the studies indicated (**Table 1**). In cases in which genes were provided as gene names or RefSeq IDs, the names were converted to their corresponding Ensembl IDs using information from the Ensembl BioMart. Half-life measurements were log-transformed after adding a pseudocount of 0.1 or 1, depending upon whether the unit of the provided half-life was measured in hours or minutes, respectively. Duplicated IDs were then averaged to compute a single half-life measurement for each gene. Five human datasets from four studies provided data as degradation rates rather than half-lives [29,38,46,49], so their transformed half-life values were negated. Finally, we generated a sparse matrix of half-lives for all genes x samples, containing missing values for genes in which the sample did not provide a measured half-life (**Additional file 2: Dataset S1**).

To evaluate the relatedness between samples, we first extracted the subset of genes in the aforementioned matrix for which ≥10 human samples (or ≥5 mouse samples) reported non-missing values. We then z-score transformed the half-lives from each sample to standardize the scale of each sample. To impute the missing values of the matrix, we used the *estim_ncpPCA* function in the *missMDA* R package to estimate the appropriate number of principal components (PC) for the *imputePCA* function, considering between 0 to 20 components for the human samples (or 0 to 10 components for the mouse samples). Finally, the data was quantile normalized using the *normalize.quantiles* function in the *preprocessCore* R package. The first PC of this imputed matrix (as computed by the *prcomp* function in R) was used as a robust cell-type-independent measurement of mRNA half-life, and computed for human and mouse species separately. For human, the samples from a single study [44] were removed prior to computing the PC because the samples from this study represented large outliers relative to all other samples (**Additional file 3: Dataset S2**).

For evolutionary comparisons between species, one-to-one human-to-mouse orthologs were acquired from the Ensembl v90 BioMart [99] by extracting the “Mouse gene stable ID” and “Mouse homology type” with respect to each human gene.

To examine sample relatedness, the first two PCs were computed using the transpose of the imputed (gene x sample) matrix, and samples were annotated and colored by their cell type of origin, study of origin, and half-life measurement technique.

### Gene annotation set

Gene annotations for protein coding genes were derived from Ensembl v83 (hg38 genome build) and v90 (mm10 genome build) for human and mouse, respectively [99]. Only protein-coding genes were carried forward for analysis. Out of all transcripts corresponding to each gene, the one with the longest ORF, followed by the longest 5′ UTR, followed by the longest 3′ UTR was chosen as the representative transcript for that gene [2,24,25].

### Basic mRNA features to predict half-life

The G/C content and lengths of each of these functional regions (i.e., 5′ UTRs, ORFs, and 3′ UTRs), intron length, and ORF exon junction density (computed as the number of exon junctions per kilobase of ORF sequence) were gathered as “basic” mRNA features associated with mRNA half-life [2,6,7]. All length-related features were transformed such that: ^^^x ← log_10_(*x* + 0.1) to reduce the right skew [2].

### Sequence features to predict mRNA half-life

All genetically encoded features are summarized in **Table 2**. Codon frequencies were extracted from ORF sequences using the *oligonucleotideFrequency* function (parameters width=3, step=3) in the *Biostrings* R package, and normalizing the counts for each codon by the sum of all codon counts for the gene. K-mer frequencies were computed similarly from each of the 5′ UTR, ORF, and 3′ UTR sequences (parameters width={1..7} depending on the size of the k-mer, step=1).

MicroRNA features were collected for mammalian-conserved miRNA families using TargetScanHuman7.2 and TargetScanMouse7.2 [24]. We computed a miRNA target binding score by negating the “cumulative weighted context++ score” for each miRNA family.

The degree of predicted binding to a number of RBPs was computed for 5′ UTR, ORF, and 3′ UTR sequences separately using SeqWeaver [68] and DeepRiPE [69]. For SeqWeaver, we generated predictions on each 50 nt window, padding the sequence with its neighboring 475 nt upstream and downstream sequence (or Ns in the case of sequences at the boundary of a region) to generate a 1000 nt input sequence. For DeepRiPE, we generated predictions on each 50 nt window, padding the sequence with its neighboring 50 nt upstream and downstream sequence (or Ns in the case of sequences at the boundary of a region) to generate a 150 nt input sequence. DeepRiPE additionally required information about the functional region of interest, which we provided according to the functional region being considered. After generating predictions for all human and mouse RBPs for each functional region and gene, we computed the average value of the binding by normalizing the sum of values for each predicted RBP to the total length of the functional region.

### Biochemical features

All biochemical features are summarized in **Table 2**. The number of CLIP peaks associated with each gene for each of 133 RBPs was downloaded from the ENCORI database [71]. eCLIP peaks were collected from the ENCORE browser as narrowPeak BED files [72]. The peaks for each RBP were intersected with gene body annotations and counted for each gene using “bedtools intersect” (parameters -s -c) [100]. PAR-CLIP peaks were downloaded from a previous study [70] and processed similarly to count the total peaks overlapping the gene body. m6A pathway (*i.e.*, a6A, m6Am, YTHDF2, METTL3, METTL14, and WTAP) CLIP peaks were collected from an assortment of previous studies [37,42,73–77], and if not already provided, the number of peaks intersecting gene bodies was computed. For all CLIP assays, genes with missing values (*i.e.*, those without an annotated peak) were considered to have zero peaks. All peak counts were transformed as such: ^^^x ← log_10_(*x* + 1) to reduce the right skew of the distributions.

Processed RIP-seq and translational efficiency data was downloaded from previous studies [36,78–80]. Merging these values with half-lives resulted in missing values; these were imputed using the *impute* function of the *imputeR* R package (parameter lmFun=“plsR”) when considered alongside the “basic” continuous features describing mRNA properties.

### Lasso regression modeling

All features being considered in a model (*i.e.*, basic, genetic, biochemical, and/or a subset within each category) and half-life values were concatenated together and z-score normalized by subtracting their respective mean values and dividing by their standard deviations. We trained a lasso regression model on each of 10 folds of the data. A lasso regression model was chosen specifically because it employs an L1 regularization penalty, which leads to the selection of the fewest features that maximally explain the data. The strength of the regularization was controlled by a single *λ* parameter, which was optimized using 10-fold CV on the entire dataset. To evaluate the usefulness of different subsets of features in improving predictive performance, we evaluated the Pearson correlation of the predictions of the lasso regression model on each of the 10 held-out folds of data, and performed paired t-tests to evaluate significant improvements in performance. To interpret the best model, we trained the lasso regression model on the full dataset and visualized the top 30 coefficients with the greatest magnitude.

### Saluki model architecture and training

We trained a hybrid convolutional and recurrent deep neural network to predict half-life from its spliced mRNA sequence with several important gene structure annotations. Following previous work on applying such models to DNA/RNA sequence, we one-hot encoded the nucleotide sequence to four input tracks. Due to the strong influence of splicing on RNA stability, we added a fifth binary track to mark the positions of exon junctions at each exon’s 5′ nucleotide. Due to the strong influence of mRNA codon composition, we added a sixth binary track to mark the beginning nucleotide of each codon, which implicitly labels the 5′ and 3′ UTRs due to their absence of codon markers. Although a nucleotide-only strategy may be preferable for mutation effect prediction, these mRNA features are important and could not easily be predicted by the model without adding substantial auxiliary training information.

Because mRNA lengths vary by orders of magnitude, we designed a model architecture to work with variable length sequences. We capped the length of an input mRNA to the model at 12,288 nt for practical purposes, finding that consideration of longer sequences led to equal performance at the cost of slower training speed. To accommodate the rare scenario in which an mRNA exceeded 12,288 nt, we truncated such cases from the 5′ end, restricting the model to access the 3′-most 12,288 nt. Conversely, for the common scenario in which an mRNA was shorter, we padded such cases with zeros at their 3′ ends, representing “N”s. We made use of the ten folds of human genes described above, and divided the mouse genes into ten folds that maintain the closest homologous genes in the same fold across species based on Ensembl annotations [99].

Our model architecture, which we refer to as Saluki, consists of a “tower” of six convolution blocks to reach a resolution for which each position represents 128 nt (Fig. 5a and **Additional file 1: Fig.** S6a). Each block includes the following operations: (i) layer normalization [101], (ii) ReLU activation, (iii) 1D convolution with kernel width 5, (iv) dropout, and (v) max pooling with width 2. Overall, the model consists of 155,521 learnable parameters. We chose layer normalization over batch normalization because most of the 3′ positions are zero padded and would confuse the batch statistics. In contrast, layer normalization is computed independently at each position and simply maintains a zero vector for the padded regions.

To make a single numeric prediction for each sequence, we must aggregate information across the variable lengths. To achieve this, we use a common recurrent neural network block called the Gated Recurrent Unit (GRU) [102]. After layer normalization and ReLU activation, the GRU runs backward from the often-padded 3′ end to the information dense 5′ end. We take the final GRU hidden representation from the most 5′ position as a summary of the entire sequence. We apply a subsequent dense block, consisting of batch normalization, ReLU, and a dense layer. Finally, we apply one more block of batch normalization, ReLU, and a final dense transformation to produce the half-life prediction.

We trained with the MSE loss function using the Adam optimizer on batches of 64 examples and learning rate 0.001, beta1 0.9, and beta2 0.98. We clipped gradients to a global norm of 0.5. We used dropout probability 0.3 throughout and added L2 regularization on all convolution, GRU, and dense layer weights with coefficient 0.001. We used skopt to optimize these hyperparameters as well as the number of channels throughout the model. The hyperparameters specified here and 64 channels achieved the greatest validation set accuracy and compose our final model. For model training, we performed early stopping after 25 epochs without improvement and took the final parameters that achieved the greatest Pearson correlation on the validation set.

We trained models using both the human and mouse data using a published approach [5], in which all parameters are shared except for two separate final dense blocks for human and mouse. During training, we iterate between interwoven batches of human and mouse genes.

Our ten folds of genes allowed us to train multiple models, in which each fold was held out as a test set (and another was held out as a validation set). Because we observed variance from training run to training run, we trained five replicate models for each held out test fold, producing a total of fifty trained parameter settings. After preliminary analyses to examine the predictions’ variance, we averaged the predictions of the five replicates per test set as an ensemble, which improves accuracy and robustness. For all downstream analyses involving Saluki predictions on third-party test sets, we averaged the predictions from all fifty models.

### *In silico* mutagenesis with Saluki

We performed in silico saturation mutagenesis (ISM) to predict the effect on mRNA half-life of all transcriptomic nucleotides. For each position, we ran three Saluki forward passes, mutating the reference nucleotide to each of the three possible alternative alleles. For each mutation, we compared the half-life prediction to the reference.

For mutations that modify stop codons, we optionally set the coding track 3′ onwards to zeros. This mode creates disproportionately large effect predictions for these mutations, which can be inconvenient for analyses focused on alternative aspects. We therefore decided not to modify the coding track downstream of the stop codon.

### Insertional motif analysis with Saluki

Using our ISM scores as input, we ran TF-MoDISco [83] on each functional region (i.e., 5′ UTR, ORF, and 3′ UTR) independently to identify the enriched motifs associated with changes in half-life. We selected several of the resulting PWMs to perform an insertional analysis, choosing the consensus sequence as a representative kmer to insert. We inserted each kmer into one of 50 evenly-divided positional bins along each functional region of a valid mRNA, replacing the reference sequence with the inserted kmer to preserve the length of the mRNA. A valid mRNA was defined as one whose 5′ UTR length was ≥100nt, ORF length was ≥500nt, and 3′ UTR length was ≥500nt. For each insertion, we recorded the predicted change in half-life relative to the corresponding wild-type mRNA. Finally, for each of the 150 positional bins, we averaged the predicted changes in half-lives across all valid mRNAs to calculate the average influence of the motif across heterogeneous sequence contexts. We performed insertional analysis identically with the 61 non-stop codons, except that each codon was inserted into the first reading frame within the ORF in which it was inserted.

## DECLARATIONS

### AVAILABILITY OF DATA AND MATERIALS

The code to reproduce the core results of this work is provided under the Apache2.0 open access license at the following links: https://github.com/vagarwal87/saluki_paper (to reproduce figures) and https://github.com/calico/basenji/tree/master/manuscripts/saluki (to train deep learning models).

## AUTHOR CONTRIBUTIONS

V.A. and D.R.K. conceived of the study. V.A. designed and performed most analyses. D.R.K. trained deep learning models. V.A. generated figures and wrote the paper with feedback from D.R.K.

## ACKNOWLEDGMENTS

We thank Han Yuan for help with TF-MoDISco analysis. We also thank Han Yuan, Divyanshi Srivastava, Nimrod Rubenstein, Charles Ledogar, and David Botstein for critical feedback.

## SUPPLEMENTARY INFORMATION

**Additional file 1: Figures S1-S7.** Supplementary figures S1 to S7.

**Additional file 2: Dataset S1.** Sparse matrices of human and mouse half-lives collected for each sample, provided along with the corresponding Ensembl gene IDs and gene names. Missing values exist for mRNAs whose half-lives were not measured or provided.

**Additional file 3: Dataset S2.** Matrices of filtered, transformed, imputed, and quantile-normalized human and mouse half-lives collected for each sample, provided along with the corresponding Ensembl gene IDs and gene names. Also provided are the aggregated half-lives for each species, computed as PC1 of each matrix.

## SUPPLEMENTARY FIGURES

**Figure S1.**
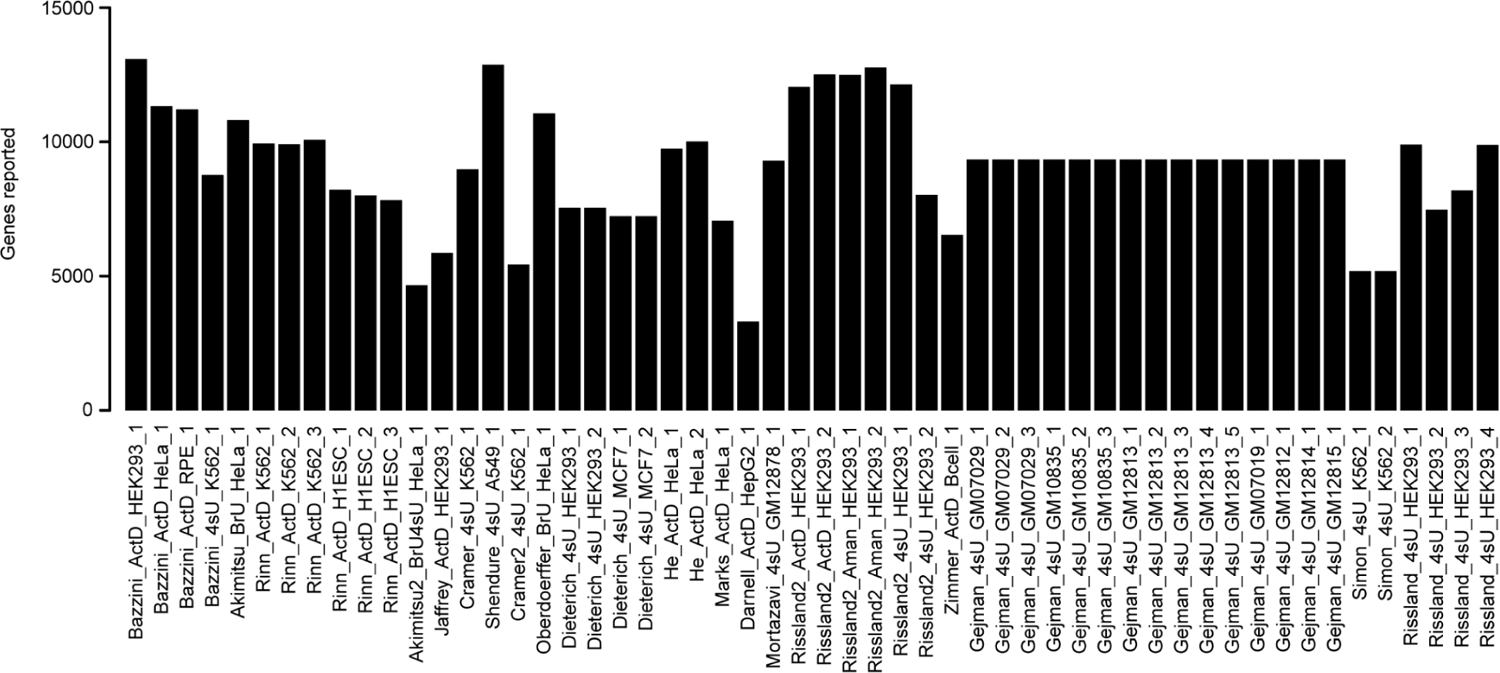
Variability in genes reported in different human half-life datasets. Barplot of the number of genes whose half-lives were reported in each of 54 human samples. Each sample ID is delimited by underscores and listed according to its study of origin (**Table 1**), measurement method, cell type, and replicate number.

**Figure S2.**
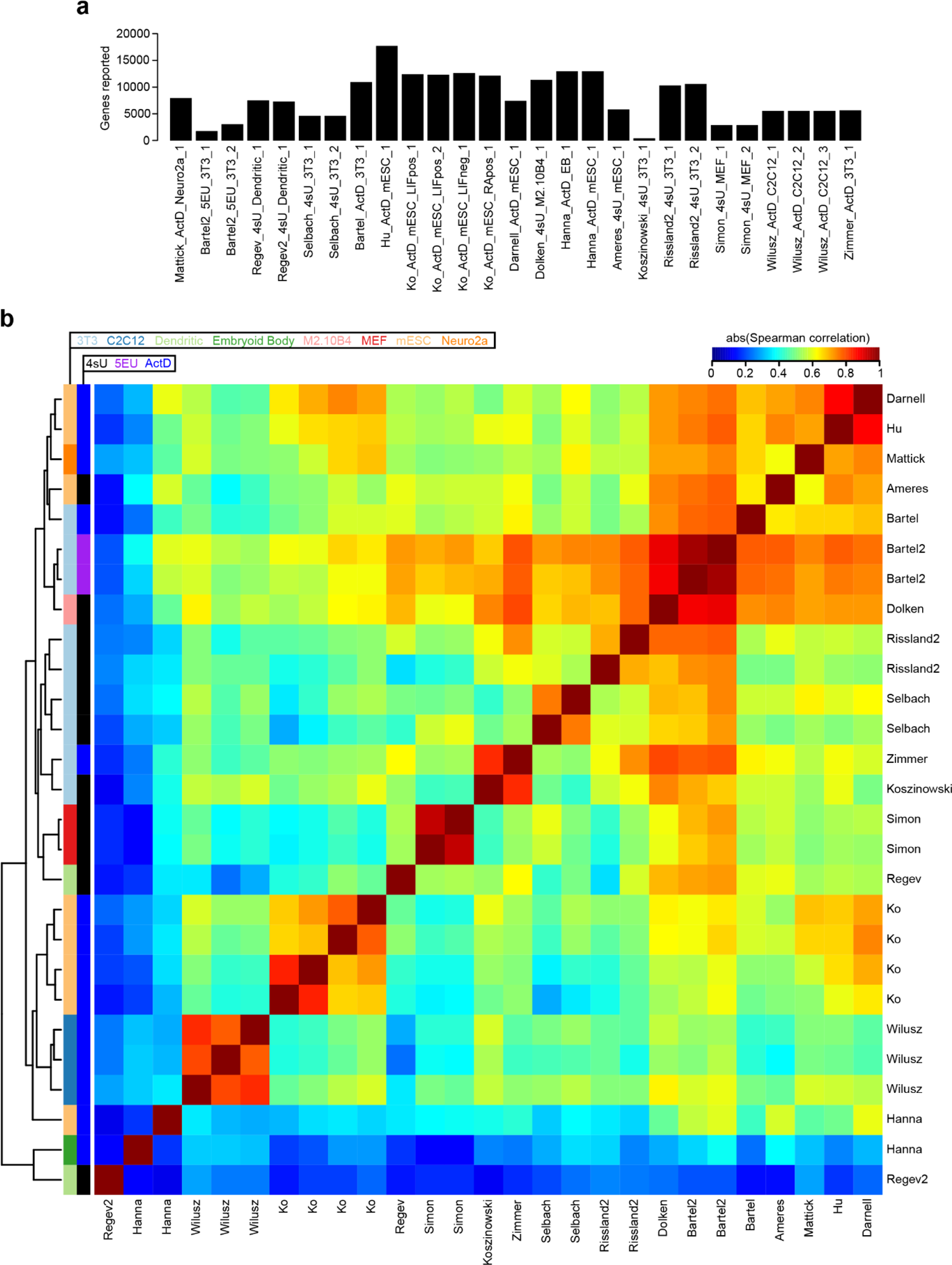
Comparison of half-lives in a compendium of mouse datasets. **a)** Barplot of the number of genes whose half-lives were reported in each of 27 mouse samples. Each sample ID is delimited by underscores and listed according to its study of origin (**Table 1**), measurement method, cell type, and replicate number. **b).** Heatmap of the absolute value of the Spearman correlations measured between half-lives derived from each pair of 27 mouse samples. Samples are clustered using hierarchical clustering according to the indicated dendrogram. Rows are labeled by the study of origin (**Table 1**) and colored by the cell type of origin and measurement approach.

**Figure S3.**
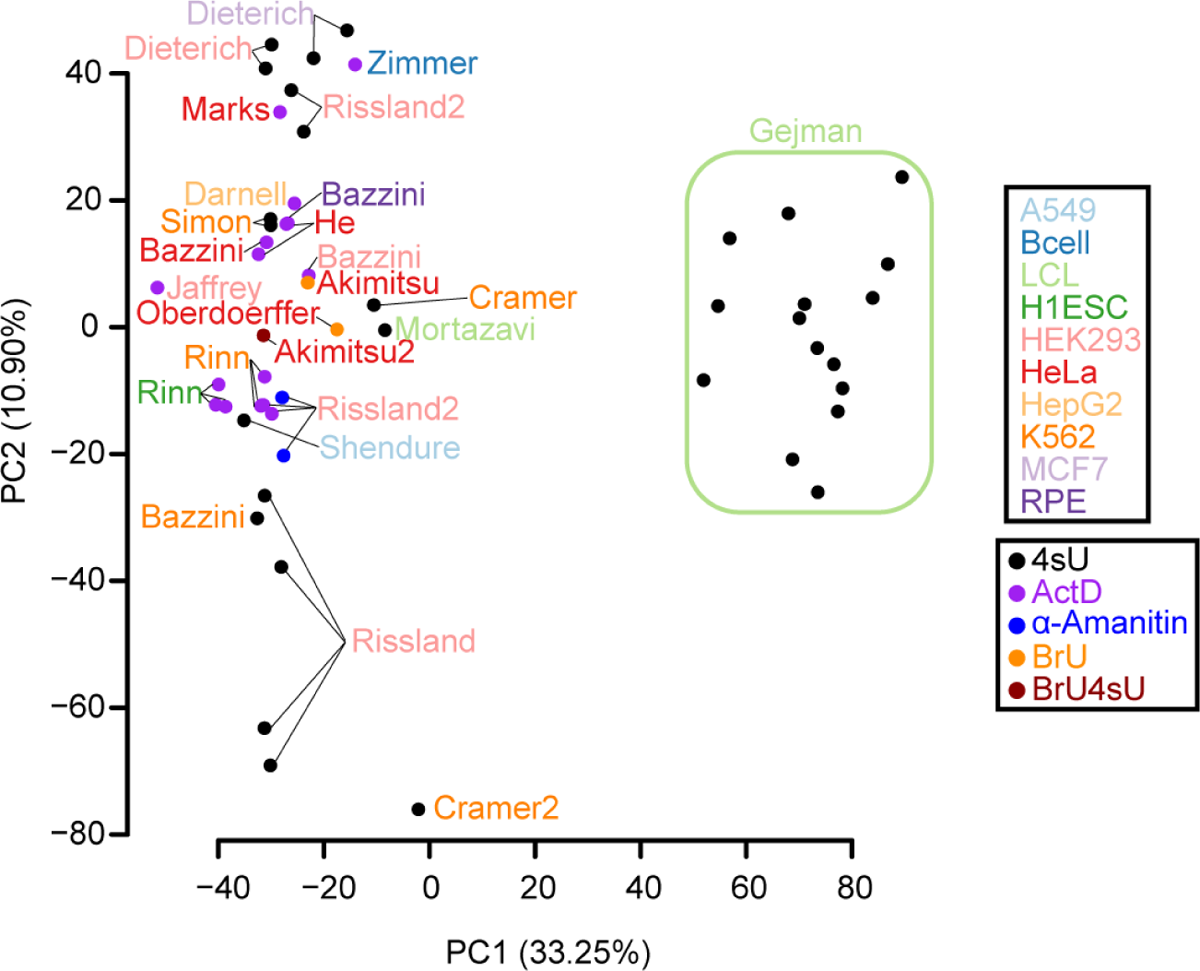
Relationship among human samples. This figure is similar to Fig. 2a, except that it shows all human samples evaluated in this study.

**Figure S4.**
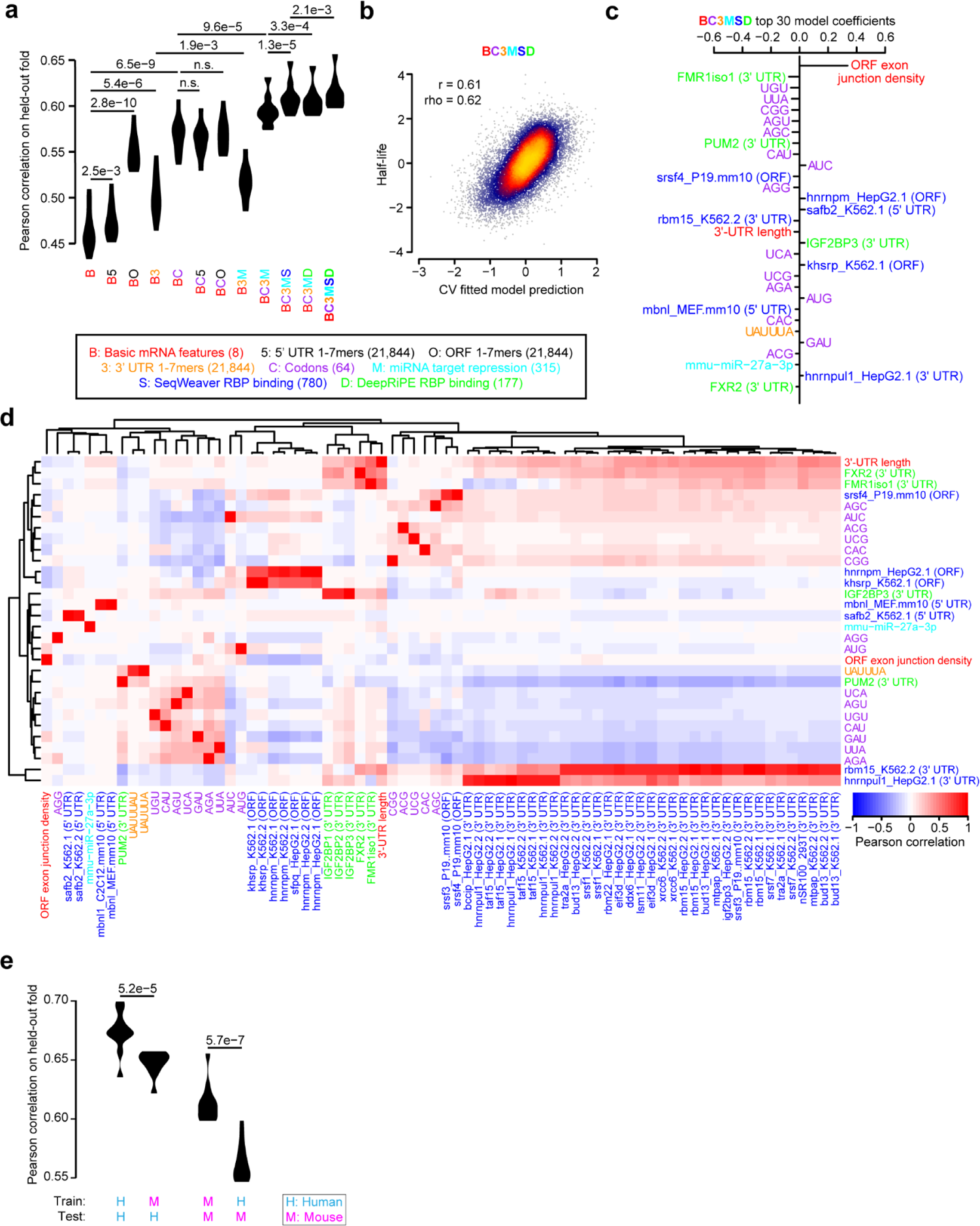
Prediction of mouse half-lives using sequence-encoded features. **a-d)** These panels are organized in the same fashion as **Fig. 3a-d**, except that they evaluate features benchmarked upon mouse data rather than human data. **e)** Performance of trained human and mouse models on test sets derived from either the same or different species. Due to the imperfect one-to-one miRNA mappings between the human and mouse, miRNA-related coefficients were excluded from the inter-species predictions. Each statistical comparison shown was evaluated with a paired t-test.

**Figure S5.**
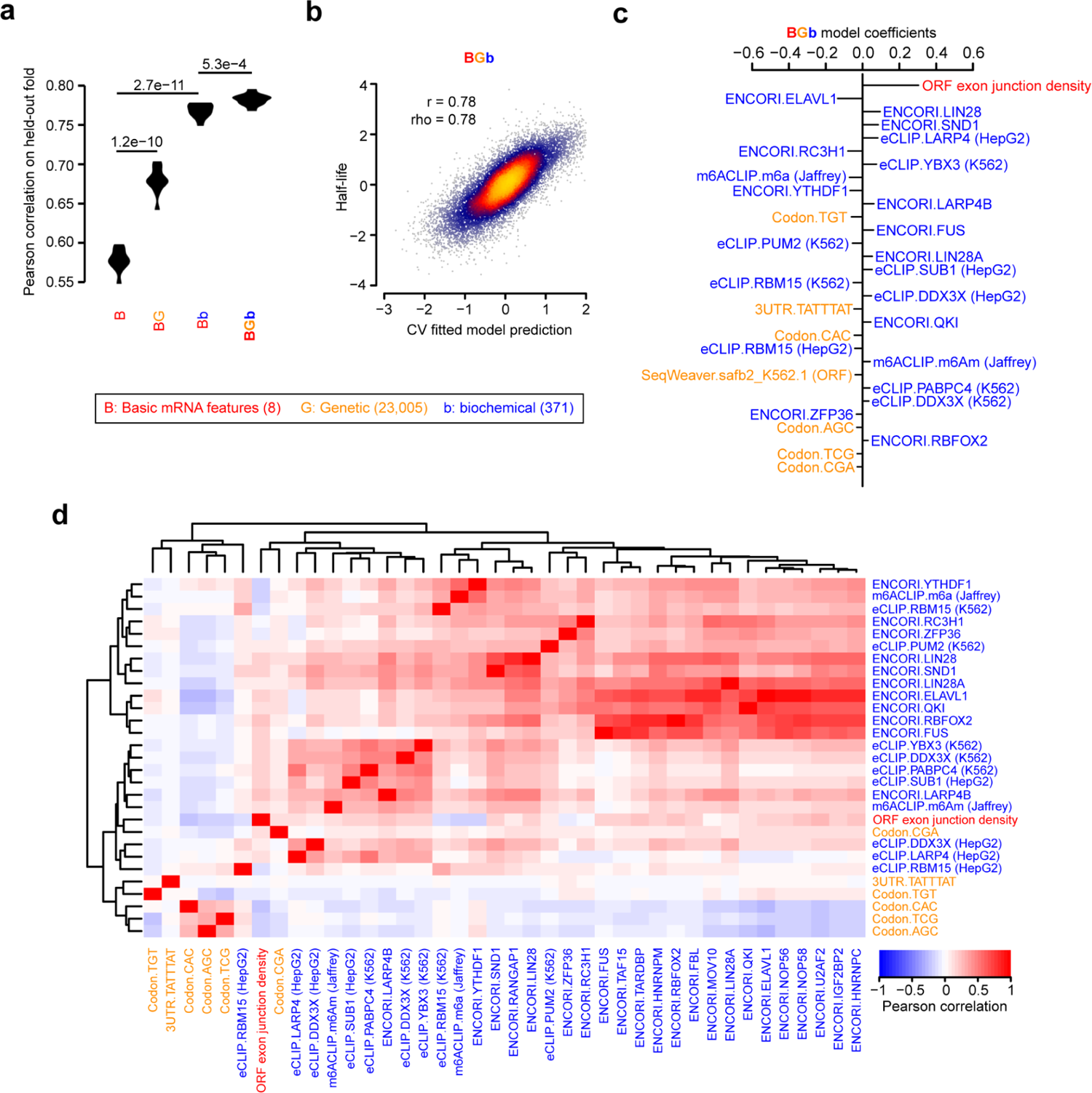
Prediction of mouse half-lives using biochemical and sequence-encoded features. This figure is organized in the same fashion as **Fig. 3-4**, except that it evaluates the subset of optimal genetic features (BC3MS model, Fig. 3), optimal biochemical features (BEeM model, Fig. 4), or a combination of both genetic and biochemical features. The features shown in panels (c-d) are described alongside their corresponding source, such as eCLIP, SeqWeaver, or codon features.

**Figure S6.**
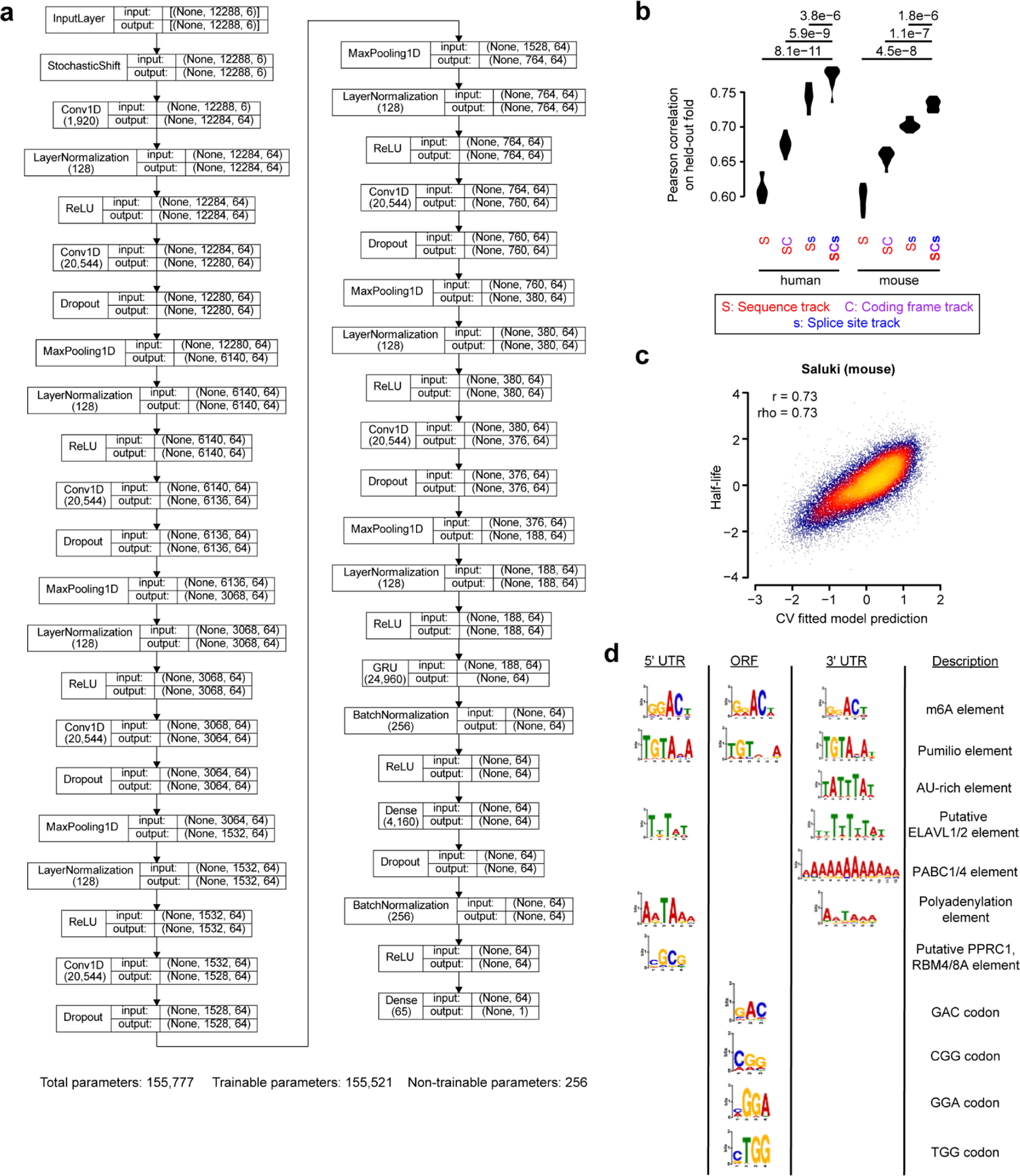
Performance and interpretation of Saluki. **a)** Complete architecture of the Saluki model. Indicated for each layer is the layer name, number of parameters in parentheses when applicable, and dimensionality of the input and output matrices. **b)** Performance of trained Saluki models on each of 10 held-out folds of data. Compared is the relative performance between pairs of models, for both human and mouse species, which iteratively consider additional input tracks. Each model is described by a code indicating the input track considered. A description of the code is provided in the key. An improvement in a more complex model relative to a simpler model was evaluated with a one-sided, paired t-test, adjusted with a Bonferroni correction to account for the total number of hypothesis tests. **c)** This panel is organized in the same fashion as Fig. 5c, except that it evaluates the performance on mouse data rather than human data. **d)** Set of enriched motifs discovered by TF-MoDISco [83] in each of the three functional regions of an mRNA.

**Figure S7.**
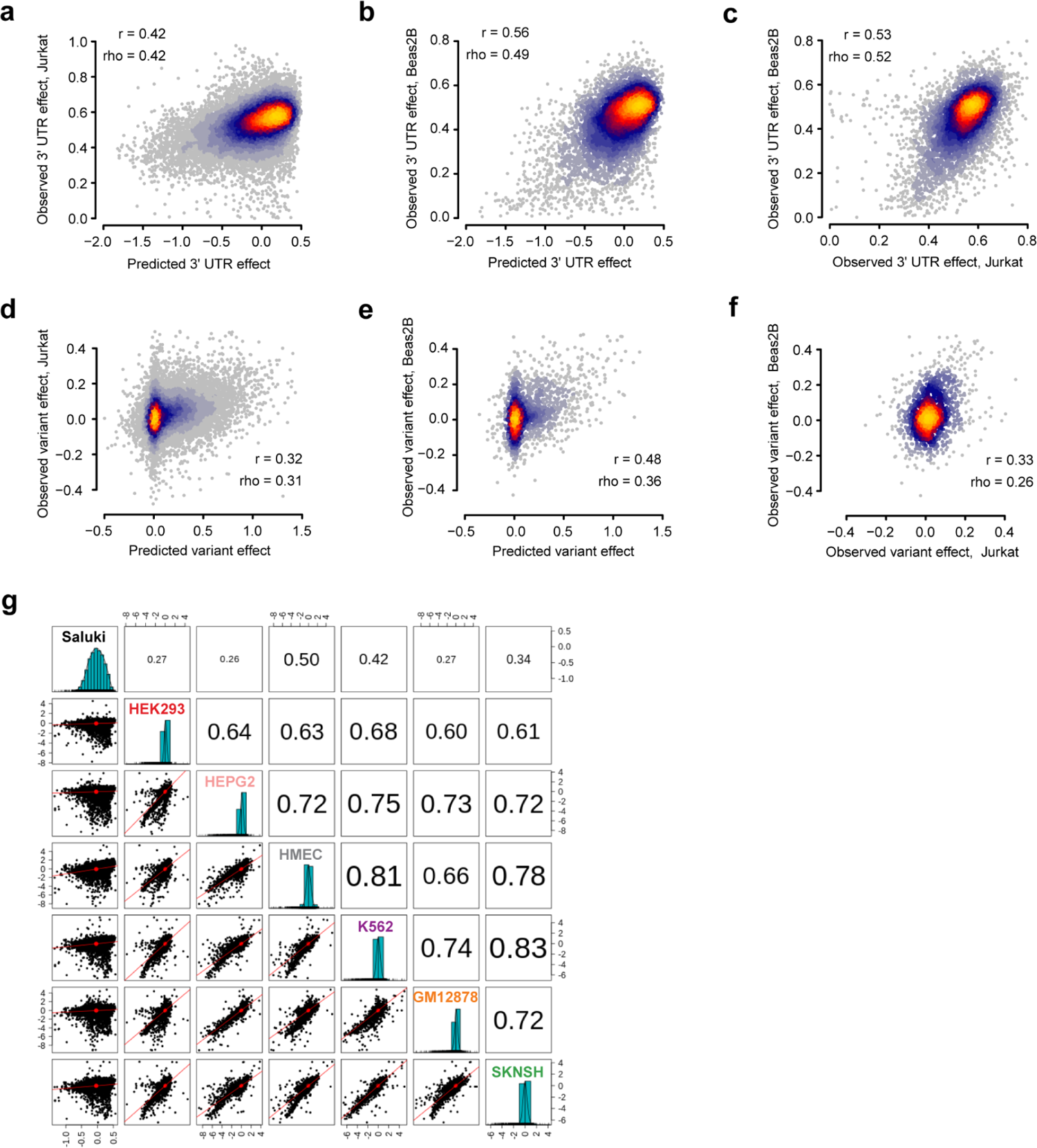
Concordance of Saluki predictions and additional functional data from massively parallel reporter assays. **a-c)** Scatter plot of the observed and predicted 3′-UTR effects, as measured in a) Jurkat cells or b) Beas2B cells, alongside c) a plot of observed 3′-UTR effects between the pair of cell types [87]. **d-f)** These panels are organized like panels (a-c), but display variant effects instead of 3′-UTR effects [87]. Panel e) is identical to that shown in Fig. 6e and reproduced here for convenience. Also indicated are the Pearson (r) and Spearman (rho) correlation values for panels (a-f). **g)** Scatter matrix displaying scatter plots corresponding to each of the 21 pairs of possible comparisons (lower diagonal elements) involving Saluki predictions as well as the measured stability effects for 3′-UTR fragments measured in each of six cell types [88]. Shown on the diagonal is a histogram of the predicted or observed RNA half-life scores. Also shown are Spearman correlation values among each pair of comparisons, with the size of the text proportional to the magnitude of the correlation coefficient (upper diagonal elements).

